# Navigated range expansion promotes migratory culling

**DOI:** 10.1101/2024.03.09.584265

**Authors:** Yi Zhang, Qingjuan Hu, Yingtong Su, Pan Chu, Ting Wei, Xuefei Li, Chenli Liu, Xiongfei Fu

## Abstract

Motile organisms can expand into new territories and increase their fitness, while nonmotile viruses usually depend on host migration to spread across long distances. In general, faster host motility facilitates virus transmission. However, recent ecological studies have also shown that animal host migration can reduce viral prevalence by removing infected individuals from the migratory group. Here, we use a bacteria-bacteriophage co-propagation system to investigate how host motility affects viral spread during range expansion. We find that phage spread during chemotaxis-driven navigated range expansion decreases as bacterial migration speed increases. Theoretical and experimental analyses show that the navigated migration leads to a spatial sorting of infected and uninfected hosts in the co-propagating front of bacteria-bacteriophage, with implications for the number of cells left behind. The preferential loss of infected cells in the co-propagating front inhibits viral spread. Further increase in host migration speed leads to a phase transition that eliminates the phage completely. These results illustrate that navigated range expansion of the host can promote the migratory culling of infectious diseases in the migration group.

**Significance Statement:** Host migration is commonly believed to accelerate the spread of infectious diseases. However, recent ecological studies suggest that migration may impede this spread. In our study, we developed a synthetic host-virus co-propagation model to explore the impact of host range expansion on the interplay between host mobility and virus spatial distribution. Our experimental and theoretical analysis revealed the spatial sorting of uninfected and infected hosts in the navigated propagating front leads to faster back diffusion of infected hosts. This self-organized structure allowed the migrating host population to eradicate the infectious disease, independent of intricate host-virus dynamics.

## Main Text

Range expansion is a process by which living species invade new territories and gain benefits such as survival(1–3), reproduction(4, 5), and resources(6, 7). Classic theory predicts that motile populations expand in space and time by growing and moving(8–10). Recent studies have shown that many organisms can use self-generated cues, known as navigated range expansion, to increase the expansion rate toward long-term settlement(11–14). However, nonmotile species, have to use other strategies to expand their territories, such as viruses, which depend on the infected hosts to spread across long distances(15–17). It is commonly assumed that host motility enhances virus transmission and viral range expansion(18–21). However, recent ecological studies have also reported that animal host migrations can reduce viral prevalence(22–25). This can occur when animals escape from infected regions (‘migratory escape’)(26, 27) or when infected animals are removed from migrating populations (‘migratory culling’)(22, 28). One of the prominent examples is that the seasonal migration of butterfly monarchs reduces the risk of infection of parasites(22, 29). These conflicting findings from epidemiological case studies suggest that the effects of host spatial expansion on viral prevalence depend on the specific conditions of the system, which require a close and quantitative examination. However, studying infectious disease dynamics and their mechanisms in field studies is difficult due to the limitations of observatory technology and the complexity of the natural system. Therefore, we developed a co-propagation system with motile bacteria, *Escherichia coli* (*E*.*coli*), and its chronic bacteriophage M13 which does not kill infected bacteria cells when producing progeny phage after infection(30), to investigate how host range expansion affects the relationship between host motility and viral spatial prevalence.

## Results

### Phage spread during the bacterial range expansion

We adopted a swimming assay of motile bacteria by inoculating a small droplet of cells at the center of a semi-solid agar plate (green dot in Fig. 1A, 2 μL containing about 10^6^ cells). The motile bacterial population grew and expanded radially. Ahead of the expanding bacterial population, we plated a small volume of nonlethal phage M13 (light pink dot in Fig. 1A, containing about 5×10^6^ phage particles). When the bacteria encountered the phage zone, they became infected and temporarily reduced their growth (Fig. S1), resulting in a visible low-density region of the infected zone compared to the uninfected zones (Materials and Methods, Fig. S2). To better identify the infected region, we integrated a red fluorescent gene Ruby into the M13 phage genome (M13-Ruby, Fig. S3A). The infection of M13 phage introduced the fluorescent gene into the bacterial cells, which expressed the red fluorescent protein. Therefore, the infected region can be detected by the fluorescent signal measurement, which coincides with the visible low-density region (Fig. 1B, Fig. S3B). Time-lapsed fluorescent images further showed the dynamics of the fan-shape infected zone, where the boundary between the infected and uninfected zones was rapidly fixed once it emerged (Supplementary Movies 1&2).

**Fig. 1.**
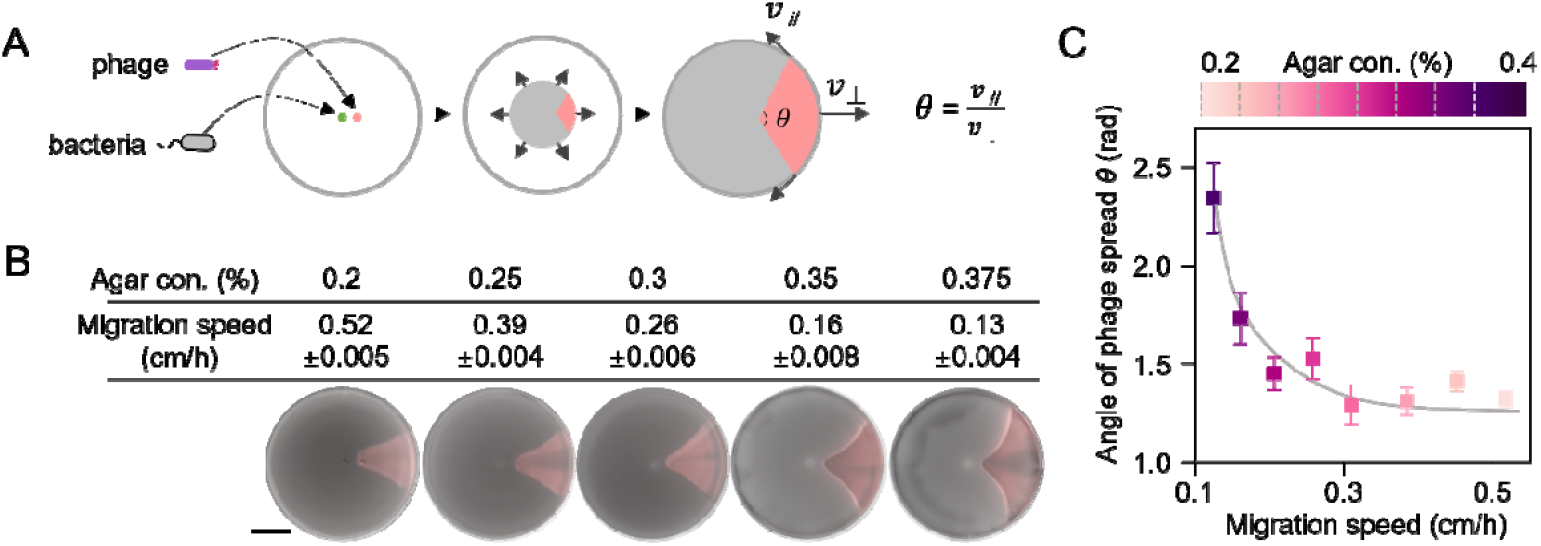
Phage spread during the bacterial navigated range expansion. (A) The co-propagation of bacteria and phage leads to a fan-shaped infected zone. (B) Typical migration speeds and fan-shaped infected zone (light pink) under different agar concentrations. *E*.*coli* FM15 was inoculated at the center of a semisolid agar plate (Materials and Methods) with varying agar concentrations, and the reporter phage M13-Ruby was inoculated 1 cm away from the center. Infected cells expressed red fluorescent protein Ruby, introduced by phage during infection. The merged pseudo-color image combines brightfield and fluorescence images. (C) Dependence of the angle of phage spread θ on bacterial migration speeds by varying agar concentrations. Strain details are given in Supplementary Table 1. Data were taken 96 hours after inoculation. Scale bar, 2 cm. Experiments in (B) were repeated independently three times. For (C), data are mean ± s.d. for n=5 biological replicates.

To understand the dynamics of bacteria-phage co-propagation, we extended a growth-expansion model that incorporates bacterial chemotaxis and M13 phage infection kinetics (Fig. 2A and SI Text). The model simulation provided a comprehensive picture about the formation dynamics of the fan-shape infected zone (Fig. S4). The infected cells produced new phages that would further infect more uninfected cells along the front of the migrating population at the same propagating speed as the host front migration speed. At the same time, the phages infected other neighboring bacteria, causing the lateral expansion of the infected region with a lateral expanding speed. The bi-directional propagation results in a fan-shaped infection region (the light pink region in Fig. 1A). The fan-shaped angle θ defines how fast the viral spread during the co-propagation with host bacteria migration, and was named the angle of phage spread. Assuming the phage infection area is approximately a sector, the angle of phage spread θ is derived as the lateral expansion speed divided by the migration speed (Fig.1A).

**Fig. 2.**
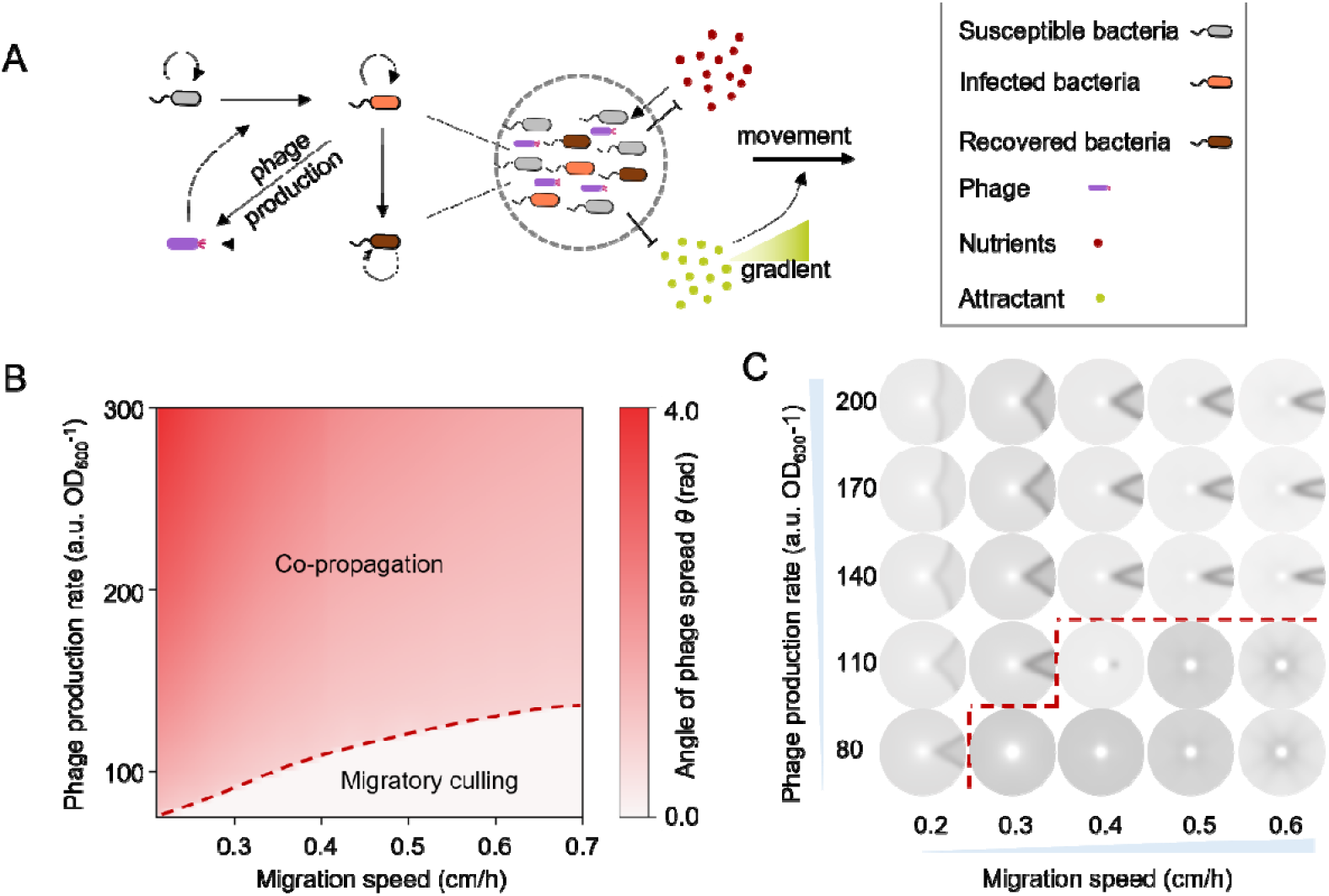
Kinetic model of the interaction between bacteria and phage. (A) Schematic illustration of population’s navigated range expansion. The M13 phage is chronic. During the interaction between bacteria and phage, the infected host cells are not killed; instead, progeny phages are produced and extruded through the cell membrane as infected cells continue to grow at a slower rate (38, 39, 56). This continues until the cells recover. Once recovered, the cells grow as fast as the susceptible ones and produce progeny phages at a much lower level as compared to freshly infected cells(38). In addition, recovered host cells cannot be re-infected by free phage particles (38). In the schematic illustration of the population’s navigated range expansion: bacterial populations navigate forward into unoccupied territories depending on the chemoattractant gradients; Bacterial populations are classified into three categories: susceptible, infected, and recovered bacteria. Infected bacteria are converted from susceptible bacteria by phage infection, and eventually become recovered bacteria. These three bacterial populations all proliferate by consuming the nutrients. Motile bacteria expand their range into unoccupied territories by diffusion and chemotaxis. Meanwhile, nonmotile phages are transmitted by their host bacteria, and their titer depends on the level of infectious progeny phage production rate by infected and recovered bacteria (Supplementary text). (B) Simulated phase diagram of bacterial-viral co-propagation. The model predicted that the angle of phage spread θ was negatively correlated with the bacteria migration speed by varying the chemotactic coefficient and positively correlated with the phage production rate. The system experienced a phase transition from co-propagation to migratory culling as further increasing migration speed at low phage production rates (red dashed line). (C) The typical fan-shaped patterns under the different migration speeds and phage production rates.

### The dependence of viral spread on the host migration speed during co-propagation

We then examined experimentally how the angle of phage spread θ depended on the host bacterial motility by varying the agar concentrations. At high agar concentrations, the probability of bacterial cells being trapped by the agar gel matrix increased, thereby reducing bacterial motility(31, 32). For the chemotactic bacterial population, which enhances their group migration by following their self-generated chemoattractant gradient (known as navigated range expansion(11)), the speed of range expansion decreased as the agar concentration increased(33). Interestingly, even though host bacterial motility was limited, the fan-shaped infected zone increased at a fixed distance from the initial infection site (Fig. 1b). We quantified the angle of phage spread θ and found it showed a negative dependence on co-propagation migration speed (Fig. 1c). However, when using a chemotaxis deficient strain (Ft-MGΔ*cheRcheB*), which performs unguided range expansion following the Fisher–Kolmogorov dynamics, in this case, the negative relation did not hold anymore (Fig. S6C-E), demonstrating that faster host motility enhanced viral spread. These results suggested a counterintuitive trend: during host navigated range expansion, an increase in host group migration speed would limit phage spread.

By varying the host diffusive motility, our model can successfully recapture the negative relation between co-propagation migration speed and viral spread (Fig. S7A). The simulation results indicated that while both co-propagation migration speed and lateral expansion speed increased with the host diffusion coefficient, the co-propagation migration speed with host chemotaxis increased at a much faster rate (Fig. S7BC).

In contrast, when considering the model without host bacterial chemotaxis (unguided range expansion, SI text), lateral expansion increased more rapidly as a function of bacteria diffusion coefficient (Fig. S5), resulting in a positive dependence of viral spread on the host motility (black line in Fig. S6D). We also confirmed that this relationship was not influenced by the geometric effect of radial expansion under the plane wave expansion (Fig. S8). Therefore, our modeling study predicted that, in the presence of host chemotaxis, increased host motility limits viral spread.

The model predicted that the co-propagation group migration speed also depended on the host chemotactic coefficient, which measures how responsive bacterial cells are to the environmental chemoattractant gradient(34). We found that the co-propagating migration speed of the host and phage increased linearly with the host chemotactic coefficient, while the lateral expansion speed stayed approximately constant (Fig. S9B, C). As a result, the angle of phage spread θ, as well as the size of the phage-infected zone, decreased significantly when the host chemotactic migration speed increased (Fig. 2B, C, Fig. S9A).

We further tested the model prediction by measuring the angle of phage spread θ with a chemotactic coefficient titratable strain (ZF1). Specifically, we deleted the native regulation of a chemoreceptor gene *tar* in the bacterial genome, and introduced a titratable control of *tar* expression by a small molecule inducer, anhydrotetracycline (aTc) (Materials and Methods, Fig. 3A). The *tar* variant strain affected the receptor gain of the cells in response to aspartate(35), but not the tumble bias and growth rate(36). The tumble bias defines the average time a cell spends on the tumbling motion, which further determines the effective diffusion coefficient of bacteria. Therefore, by titrating different inducer aTc concentrations, we could manipulate the bacterial chemotactic migration speed (Fig. 3B, Fig. S10A), but not the effective diffusion coefficient of bacteria(36). Under different host migration speeds, we measured the angle of phage spread θ and found it decreased as the migration speed increased (Fig. 3B, Fig. S10B&10C). The experimental results confirmed our model prediction that virus spread is hindered by the host chemotactic migration.

**Fig. 3.**
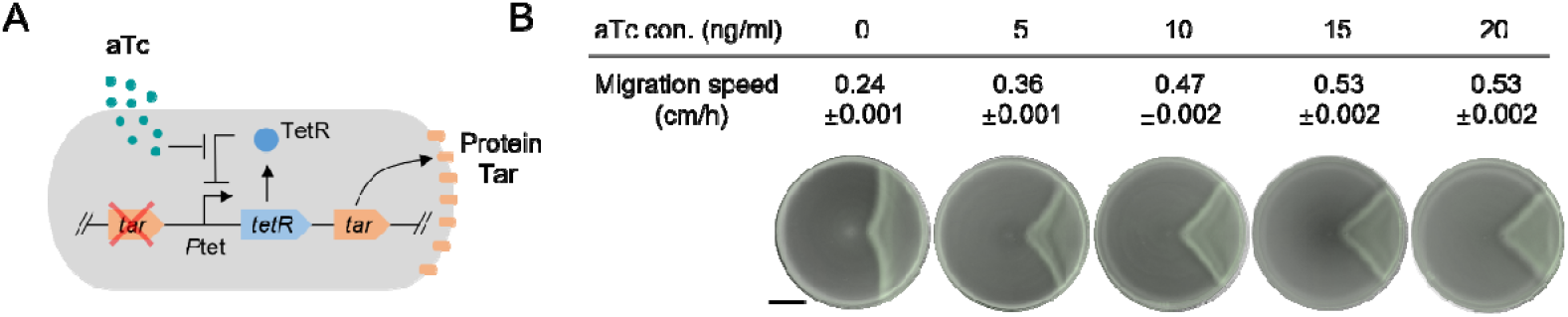
Phage spread under different co-propagating migration speeds by titration of host bacteria chemotactic abilities. (A) Design of the chemotactic ability titratable strain ZF1 by inducible expression of chemotactic receptor protein Tar (Materials and Methods). (B) Typical migration speeds and fan-shaped patterns under different aTc concentrations (Materials and Methods). Data were taken 30 h after inoculation. Scale bar, 2 cm. Data are presented as mean ± s.d. for n=3 biological replicates.

### The migratory culling of phages during co-propagation

Another important prediction of our model is that co-propagation of host bacteria and phage is not always sustained during the host navigated range expansion. As we decreased the phage production rate in the model, it resulted in a slower lateral expansion of the infected zone, limiting the angle of phage spread θ and the viral spread. At very low phage production rate, the angle of phage spread θ became small. If we further increased the group migration speed, the size of the infected zone shrank until a sudden drop to almost zero, where infection only occurred near the initial spot of phage (Fig. S11). This indicated that the co-propagation of host bacteria and phage could not be maintained and the infected cells as well as the phages were culled from the propagating front. In other words, the system underwent a phase transition from co-propagation to migratory culling (Fig. 2B&2C).

To experimentally test the phase transition of migratory culling, we engineered both the M13 phage and its bacterial host to enable the titration of the phage production rate after infection. Specifically, we first deleted the *gIII* gene from the M13 phage genome and replaced by a T7 *rnap* gene (M13SP)(37, 38). The native GIII protein is essential for the M13 phage into a host bacteria, and deletion of GIII protein prevents full escape from the infected host(38). In other words, the variant phage M13SP cannot be produced by wild-type infected cells, as it lacks *gIII* gene(39, 40). We then incorporated the *gIII* gene into bacterial host cells under a T7 promoter (strain FTT7, Materials and Methods). When the M13SP infects the engineered host FTT7, the bacteria cells express the T7 RNAP whose gene is brought by the phage, which then activates the expression of GIII protein and restores the complete production of M13SP phage. Therefore, the variant phage M13SP infects the engineered strain FTT7 and generates a fan-shaped infection zone during the co-propagation range expansion, and the dependence of the angle of phage spread θ on the host migration speed is similar to the wild-type system (Fig. 3B and Fig. 4B).

**Fig. 4.**
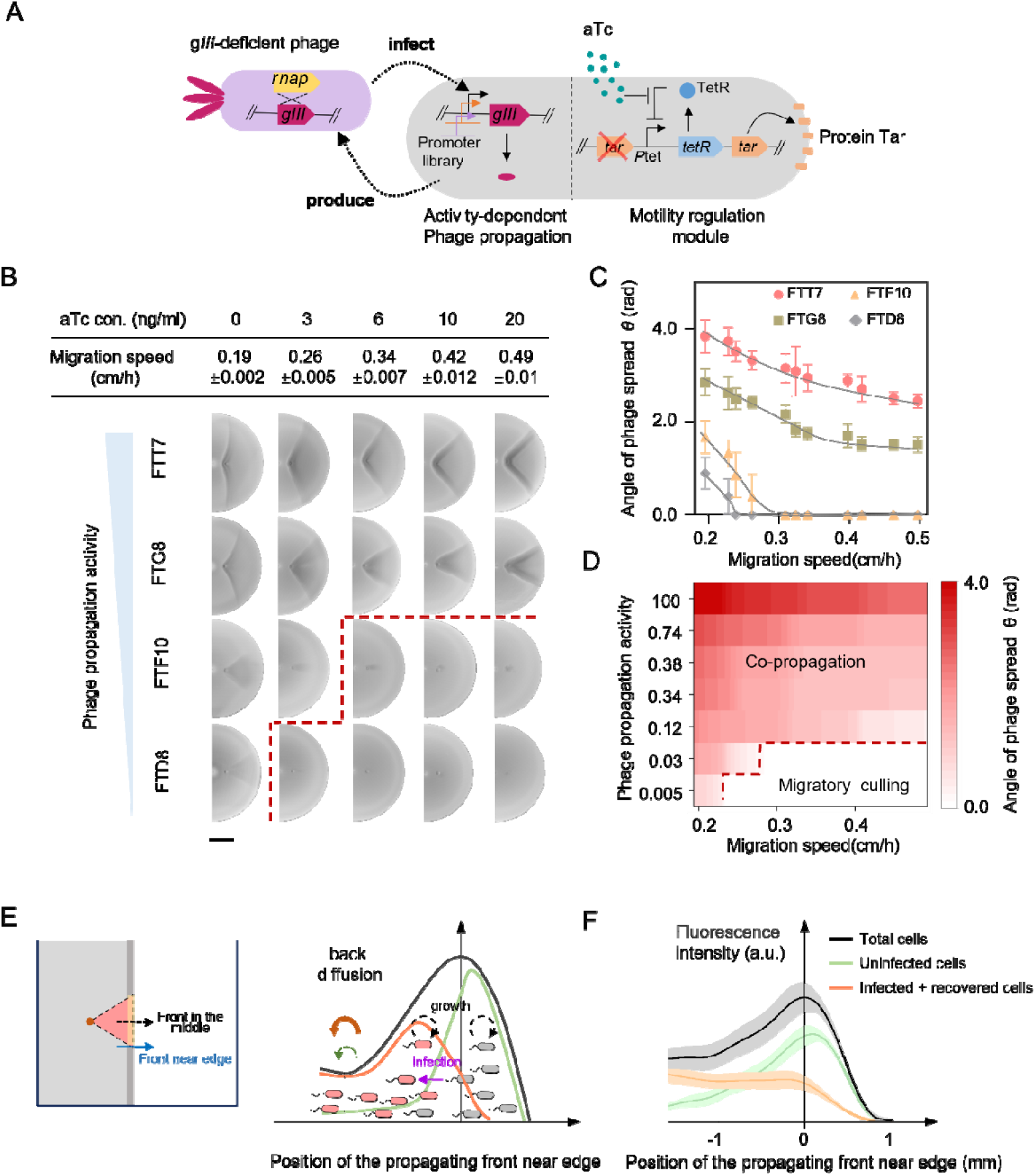
The phase transition and the spatial sorting mechanism of migratory culling. (A) Schematic design of the chemotactic titratable strain ZF1 coupling T7 RNAP activity with the expression of *gIII*. The host bacteria carry a motility regulation module and an activity-dependent phage propagation module, which are harbored by chromosome and the accessory plasmid, respectively (Materials and Methods). (B) The typical migration speeds and fan-shaped patterns for the typical strains (FTT7, FTG8, FTF10, and FTD8) under different aTc concentrations. (C) Detailed relationship between the angle of phage spread θ and the migration speed in (B). (D) Phase transition of migratory culling by seven strains with different phage propagation activities (The detailed sequences of T7 promoter variants are shown in Supplementary Table S2) under different aTc concentrations (Materials and Methods). The angle of phage spread θ was negatively correlated with the bacteria migration speed and positively correlated with the phage production rate. Migratory culling occurred at higher migration speeds and lower phage production rates. (E) Illustration of the spatial sorting mechanism for migratory culling. Along the leading propagating front near the edge between the infected and uninfected zone (as indicated by the blue arrow in the left panel), the simulated density profiles (Fig. S14) showed a spatial sorting structure of the infective cells (infected + recovered bacteria, orange line) and the susceptible cells (green line), leading to smaller back diffusion for the susceptible cell (green arrow) compared to infective cells (orange arrow). (F) Co-propagation profiles (right panel) of ZF1-RFP (a plasmid carrying red fluorescent gene rfp was inserted into the strain ZF1, Materials and Method) and phage M13-GFP (a *gfp* gene was integrated into phage genome, Materials and Methods) demonstrated the spatial sorting structure experimentally. The profiles of total (black) and infective bacteria (infected + recovered bacteria, green) were directly quantified the red and green fluorescent signals respectively, while the profile of susceptible bacteria (orange) was calculated by the other two profiles. The shaded areas represent the error bar. Data are presented as mean ± s.d. for n=4 biological replicates.

An important advantage of this engineered phage-bacteria system is that we can generate a series of engineered strains with T7 promoter variants that have different T7 RNAP binding affinities. Here, we used a library of T7 promoter variants that we had previously characterized in our study(38)(Table S2, Fig. S12). The weak T7 RNAP binding affinity, e.g. FTG8 strain, reduces the GIII protein expression(41), resulting in a smaller fan-shaped zone (Fig. 4B). Increasing the host migration speed further would lead to a much smaller fan-shaped infected zone. However, for variant strains such as FTF10 and FTD8, whose promoter expression activity is much smaller than that of wild type T7 promoter, the fan-shaped infected zone only exists at the slow host migration speed. When the host migration speed becomes large, the infected zone is confined to a small spot (Fig. 4B). Away from the initial drop of phage, we verified that no infected bacteria or phage particles were carried, suggesting they were culled from the front propagation. We then systematically measured the angle of phage spread θ for a series of variant strains at different host migration speeds (Fig. 4C&D). The phase transition from co-propagation to migratory culling is clearly observed at low promoter expression activity and fast host migration speed (Fig. 4D), demonstrating the occurrence of migratory culling.

### The spatial sorting mechanism of migratory culling

To better understand the underlying mechanism of migratory culling, we examined the simulated dynamics of the co-propagation front. During navigated range expansion, the leading front of the host population exhibited a propagating bulge, resulting from the balance between local cell growth and back diffusion (leakage of cells out of the bulge due to random movement)(3, 11). Due to phage infection, along the middle radial axis (the black dashed line in Fig. 4e), the propagating front of the host population rapidly became infected once it crossed the initial phage spot, with the phage profile generally following that of the infected cells (Fig. S13). However, simulation results showed the coexistence of uninfected and infected cells near the edge of the infected zone (along the blue arrow in Fig. 4E, Fig. S14). These cells, traveling along with phages, formed a co-propagation front (Fig. 4E, Fig. S14). Within this co-propagation front, the profile of infected cells was determined by three major factors: continuous conversion from uninfected cells due to phage infection, self-reproduction, and reduction flux caused by back diffusion. As the immotile phage always lagged behind the motile host, infection was most active behind the front of the uninfected cells. This resulted in a spatial sorting structure where the uninfected cells were located at the leading edge of the leading propagation front, while infected cells were at the rear (Fig. 4E, Fig. S14).

To validate this prediction of spatial sorting, we experimentally measured the profiles of uninfected and infected cells. We labeled the host cells with a red fluorescent gene *rfp* (ZF1-RFP) and integrated a *gfp* gene into the phage genome (M13-GFP). This allowed us to quantify the total cell density profile of the co-propagating front using red fluorescent signals (Fig. 4F, black curve, see Materials and Methods). Phage infection introduced the *gfp* gene into infected cells, enabling us to quantify the infected profile (the sum of infected and recovered cells) via green fluorescent signals (Fig. 4F, orange curve). The difference between these two provided the profile for uninfected (susceptible) cells (Fig. 4F, green curve). The experimental results clearly showed the spatial difference between uninfected and infected cells at the propagating front near the edge, which is consistent with the model prediction.

Consequently, the spatial sorting structure of the two subpopulations in the leading propagation front indicated that the back diffusion of the infected cells was larger than that of the uninfected cells. Without the conversion from uninfected cells by phage infection, the infected cells could not keep up with the uninfected migration front. In the presence of moderate phage infection (e.g. low phage production by infected cells), the increase in migration speed leads to stronger total back diffusion(11, 13), but the major increment came from the increase in the back diffusion of infected cells. Increasing the migration speed further or reducing the phage infection would make the back diffusion larger than the sum of the self-growth of infected cells and conversion rate from uninfected cells, causing the infected cells to be culled from the migration front. This resulted in a transition from co-propagation to migratory culling of infected bacterial host and phage.

## Discussion

The understanding of the interplay of host migration and viral spread has been developed with the advent of global change and advances in genetics, but direct experimental tools are still lacking (13, 22, 27). Here, we developed a synthetic host-viral copropagating system to investigate how host motility affected viral spread during the host’s navigated range expansion. We found a counterintuitive phenomenon that faster host chemotaxis-driven range expansion inhibited viral range expansion and resulted in a phase transition from co-propagation to migratory culling of phage.

Although migratory culling has been increasingly recognized in natural migratory animals and its role in regulating viral transmission dynamics, there is still a lack of agreement on how often and how much it may occur(21, 42). It is thought to depend on the extent to which infection affects the host’s physiological and behavioral traits(43). Our finding revealed that the spatial sorting of uninfected and infected hosts in the navigated propagation front resulted in a faster back diffusion of infected hosts, enabling their elimination. Even without the temporary growth reduction during infection, this mechanism still facilitated the migratory culling (Fig. S15). This quantitative understanding does not require complex infection-induced changes to host migrations, which has potential implications for controlling the spread of infectious diseases by altering the host motility or the virus production rate.

Range expansion can significantly alter evolutionary dynamics through multiple factors, e.g. genetic drift(44, 45), selection pressures(46, 47), founder effects(48, 49), and cooperation collapse(50, 51). Long-term co-propagation of host and virus can lead to co-evolution, impacting the genetic diversity and adaptation of both host and virus populations(15, 16). In addition, our previous study utilized the bacteria-phage co-propagating system to develop a spatial phage-assisted continuous evolution system, which revealed the evolutionary process during co-propagation was accelerated compared to a fixed niche(38). Therefore, the inhibition of viral spread by host migration discovered in this study would further help to improve the directed evolution method and provide new insights into host-viral co-evolution dynamics.

## Materials and Methods

### Media and growth conditions

The Luria-Bertani (LB) medium contained 10 g tryptone, 5 g yeast extract, and 5 g NaCl per liter. The defective rich defined medium (D-RDM) used in this study was based on the Neidhardt’s lab recipe(52) and modified: 1× MOPS mixture, 0.25× ACGU, 1× defective amino acid mixture (a mixture of 17 amino acid excluding asparagine, aspartic acid and serine, named as 1×AA), 1.32 mM K_2_HPO4, 100 μM aspartic acid(K salt, 0 μM only for the host unguided range expansion experiment) and 0.4% (w/v) glucose. Mops salts and ACGU were prepared to be 10× MOPS and 10× ACGU stocks solution. The defective amino acid mixture 1×AA was prepared to be 5×AA stocks solution (in 1 L): 356.4 mg alanine, 5478.2 mg arginine (HCL), 87.8 mg cysteine (HCL), 611.5 mg glutamic acid (K salt), 438.6 mg glutamine, 300.3 mg glycine, 209.6 mg histidine (HCL H_2_O), 262.4 mg isoleucine, 524.8 mg leucine, 365.4 mg lysine, 149.2 mg methionine, 330.4 mg phenylalanine, 230.2 mg proline, 238.2 mg threonine, 102.1 mg tryptophane, 181.2 mg tyrosine, 351.6 mg valine. All the medium in this study was buffered to pH 7.0 with 0.1 M HEPES (pH 7.0).

To prepare semi-solid plates, the bacto-agar (BD,214010) was added to the growth medium, and the agar concentration varied from 0.2% to 0.4% (w/v). If required, inducer aTc was also added to the medium and its concentration varied from 0 to 20 ng/mL. Then, 10 mL of the above medium supplemented was poured into a 90-mm Petri dish and allowed to harden at room temperature for 60 min in a light-proof box. In addition, the medium was supplemented with chloramphenicol (25 μg/mL), spectinomycin (50 μg/mL), tetracycline (10 μg/mL), kanamycin (10 μg/mL) and ampicillin (20 μg/mL). All experiments were executed at 37°C unless otherwise specified.

The culture condition of Fig. 1 and Fig. S8 (FM15 with different agar concentrations): D-RMD medium + 100 μM aspartate + 0.2%∼0.4% agar concentration + 10 μg/mL tetracycline;

The culture condition of Fig. S6 (Ft-MGΔ*cheRcheB* with different agar concentrations): D-RMD medium + 0.2%∼0.4% agar concentration + 10 μg/mL tetracycline;

The culture condition of Fig. 3 and Fig. S10 (ZF1 with different aTc concentrations): D-RMD medium + 100 μM aspartate +0 ∼ 20 ng/mL aTc + 0.2% agar concentration + 10 μg/mL kanamycin + 20 μg/mL ampicillin;

The culture condition of Fig. 4B,C,D, and Fig. S20 (Strains of the FT series, the chemotactic titratable strain ZF1 coupling T7 RNAP activity with the expression of *gIII*: FTT7, FTA1, FTD5, FTD8, FTD9, FTF10, FTG8): D-RMD medium + 100 μM aspartate +0 ∼ 20 ng/mLaTc + 0.25% agar concentration+ 10 μg/mL kanamycin + 50 μg/mL spectinomycin;

The culture condition of Fig. 4E (ZF1-RFP): D-RMD medium + 200 μM aspartate + 20 ng/mL aTc + 0.25% agar concentration + 10 μg/mL kanamycin;

### Strains and phages construction

The strains and plasmids used in this study are listed in Supplementary Table S1. The *E*.*coli* CLM strain was provided by Dr Liu(3) and all other strains in this study, except *E. coli* K12 strain ER2738, were derived from it. The strain FM15 was a conjugation of the *E. coli* CLM and the *E. coli* K12 ER2738 (NEB, F plasmid with tetracycline-resistant provider) and was provided by Dr Liu(38). Based on the understanding of the molecular mechanism about chemotactic signal transduction of *E*.*coli*, we constructed a chemotaxis deficient *E*.*coli* strain Ft-MGΔ*cheRcheB* by the following steps: (i) we constructed a chemotaxis deficient *E*.*coli* MGΔ*cheRcheB* by knocking the chemotactic gene *cheR* and *cheB* of *E*.*coli* CLM; (ii) we transferred the psim5 plasmid with chloramphenicol gene into the strain MGΔ*cheRcheB* and obtained the strain MGΔ*cheRcheB*-psim5 (cultured in 30°C); (iii) we conjugated the recipient strain MGΔ*cheRcheB*-psim5 with the donor strain FM15 (F plasmid provider) and got a strain Ft-MG Δ *cheRcheB*-psim5; (iv) we removed the psim5 plasmid by culturing in 37°C and obtained the strain Ft-MGΔ*cheRcheB*. The tar-titratable strain MGT was constructed as follows: (i) the strain CLM(Δ*tar*) was obtained by knocking the chemotactic receptor protein Tar gene of the strain CLM. (ii) the bla:P_tet_-*tetR*-*tar* feedback loop was amplified and inserted into the strain CLM(Δ*tar*) chromosomal attB site utilizing λRed homologous recombination, then we obtained the strain MGT. The strain FkP was constructed by replacing the *tetA-PtetA/tetR-tetR* cassette of F plasmid in the strain Er2738 with the kanamycin-resistance gene. The tar-titratable strain ZF1 was a conjugation of the strain MGT and the strain Fkp (F plasmid with Kan resistant provider). The strain FTT7 was constructed by electroporating the plasmid AP-T7 (T7 RNAP-dependent accessory plasmid contain M13 phage gene *gIII* with a wild type T7 promoter, gift from Dr Chenli Liu(38)) into the strain ZF1, and the similar strain FTA1/FTD5/FTD8/FTD9/FTF10/FTG8 was constructed by electroporating the plasmid AP-A1/ AP-D5/ AP-D8/ AP-D9/ AP-F10/ AP-G8 (The variants of plasmid AP-T7 with T7 promoter variants, and the details of the promoter sequence are shown in Appendix Table S1, gifts from Dr Liu(38)) into the strain ZF1. To better capture the location difference of the infected and uninfected cells in the co-propagating front, we constructed the strain ZF1-RFP, in which a red fluorescence plasmid *RFP* with chloramphenicol gene was inserted into the strain ZF1.

The phage M13 used in this study was a gift from Dr Liu(38) and others were its variants. To better characterize the infectious state of the bacteria, the coding sequence of green/red fluorescence protein gene was inserted into the genome of M13 and the strain was designated as M13-GFP, M13-RFP and M13-Ruby. The phage M13SP was also provided by Dr Liu(38), in which gene *gIII* of the M13 genome was deleted and replaced by a T7 RNP with a downstream yellow fluorescence protein gene. All phages were verified by sequencing.

### Conjugation of F plasmid

The conjugation of F plasmid is based on the Barrick laboratory’s recipe(53), and the detailed protocol is as follow: (i) grow overnight cultures of donor and recipient strains in the presence of diaminopimelic acid (DAP, 0.3 mM) and appropriate antibiotic. (ii) gently spin down 1 mL culture (∼6000 rpm for 3 minutes) and wash donor and recipient cells in PBS, then repeat and resuspend in 500 µL PBS. (iii) measure cell density and combine 1:1 ratio of donor and recipient cells in micro centrifuge tube (100 µL:100 µL). (iv) plate 200 µL of mixture onto non-selective LB medium plate containing DAP, and then incubate conjugation plate overnight. (v) scrape up the conjugation mixture into a micro centrifuge tube with 1 mL PBS, vortex and gently spin down and repeat, then resuspend conjugation mixture in 1 mL of PBS and plate 100 µL of this mixture and 100 µL of a 10-fold dilution onto selective plates. (vi) pick single colonies and culture in selective media and confirm the strain via PCR amplification of the target strains.

### Growth curve measurement

The growth curves of uninfected and infected bacteria were measured in a 250-mLflask with 50 mL corresponding growth medium at 37°C, 150 rpm. The general procedure was as follows. First, the isolated bacteria from −80°C stock was streaked onto the agar plate with LB medium and cultured at 37°C overnight. Second, 3-5 single colonies were picked and inoculated into 14 mL round-bottom test tubes containing 2 mL LB medium and cultured overnight in a shaker (220 rpm, Shanghai Zhichu Instrument) as the seed-culture procedure. Third, the overnight seed-cultures were diluted into 2 mL D-RDM medium by a ratio of 1:100 and grown to log-phase the next morning; when the diluted cultures reached OD_600_ 0.2∼0.3, the diluting step was repeated. Fourth, 5 mL cultures (OD_600_ around 0.2∼0.3) were added into a 250-mL flask with 45 mL prewarmed D-RDM medium and were cultured in a water-bath shaker (150 rpm, Shanghai Zhichu Instrument). When the diluted cultures reached an OD_600_ of 0.2, 2 mL bacteria culture was added to 48 mL prewarmed D-RDM medium with or without phage. The medium was then incubated and measured. The samples were taken every 10 min for measurement of OD_600_ by using spectrophotometer reader until bacterial cells entered the stationary phase. For phage growth assay, one-milliliter samples were extracted at the same time points and filtered through a 0.22 μm pore size PES syringe filter to remove bacterium. Aliquots for the time series were then stored at -20°C until tittering. The growth curve was illustrated in Fig. S1.

### Phage propagation and tittering

The strain FM15 was served as host bacteria for propagating and tittering M13 phage. Cells used for phage propagation were cultured in 20 mL LB medium until they reached OD_600_= 0.3∼0.4 and then infected with 10^9^ pfu of phage. The bacteria-phage mixture was incubated overnight at 37°C. Cell debris was removed by centrifugation at 5000 rpm for 10 min and filtration. The fresh supernatant containing the revived phage was collected and stored at -20°C for up to several months. To determine the phage titer, we employed the double agar overlay plaque assay as described in previous work(54). The bottom agar for plates and soft agar for overlayers were LB broth containing 1.5% and 0.4% Bacto agar, respectively. Serial ten-fold dilutions of the phage stock and the underlay agar plates were prepared in advance. We transferred 10 μL of the selected dilution of phages to a tube of 3 mL warm overlay medium, immediately added 100 μL culture of the host bacterium (OD_600_ around 0.3∼0.4), mixed and poured the contents over the surface of a dried and labeled underlay plate. The overlays were allowed to harden for 30 min and then incubated at 37°C overnight. The following day, we counted plaques on plates with 30– 300 plaques and defeminated the titer of the original phage stock by using the following calculation: Number of plaques×100×reciprocal of counted dilution = pfu/mL.

### Expansion experiment procedures

First, the isolated bacteria from −80°C stock was streaked onto the agar plate with LB medium and cultured at 37°C overnight. 3-5 single colonies were picked and inoculated into 14 mL round-bottom test tubes containing 2 mL LB medium and cultured overnight in a shaker (220 rpm., Shanghai Zhichu Instrument) as the seed-culture procedure. Second, the overnight seed-cultures were diluted into 2 mL D-RDM medium by a ratio of 1:100 and grown to log-phase the next morning, and then were further diluted in the same way when the diluted cultures reached OD_600_ 0.2∼0.3. In the experiment of measuring the host’s range expansion speed, bacteria were then cultured to the mid-log phase (OD_600_ was around 0.2∼0.3), which was inoculated at the line 2 cm away from the center of a semi-solid agar plate using a 75-mm glass slide for the plane-wave range expansion. Alternatively, 2 μL of the strain was inoculated at the center of a semi-solid agar plate for the normal range expansion. In the virus infection experiment procedures, 2 μL M13 phages with different concentrations were inoculated 1 cm away from the bacteria position, and then incubated at 37°C for several hours until the bacteria occupied the whole plate. For the experiment of measuring bacteria expansion speed, only the bacteria strain was inoculated and incubated at 37°C.

### Bacteria expansion speed measurement

The semi-solid agar plate was illuminated from below by a circular white LED array with a lightbox as described previously(55) and imaged at 1 h or 2 h intervals using a Canon EOS 600D digital camera. Images were analyzed using ImageJ. For the normal range expansion, a circle was fitted to the intensity maximum in each image and the area (A) of the fitted circle was determined. The radius (r) of the colony was calculated as 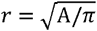. The maximum expansion speed was calculated using a linear fit over a sliding window of at least five time points, with the requirement that the fit has an R^2^ greater than 0.99. For the plane-wave range expansion, an image analysis script using MATLAB was written to find the front peak position of the images at different times and then we got the mean migration speed by a linear fitting.

### Calculation of the angle of phage spread *θ*

When the semi-solid agar plate was occupied by bacteria, brightfield images of the plates were captured by the image device in the above Methods and fluorescence images were captured using the UVP CHEMSTUDIO™ TOUCH 815 IMAGERS (Analytik Jena, US). Virus spread intensity can be represented by the angle θ of the fan-shaped pattern and can be determined as follows. Using ImageJ analysis to fluorescence images, the phage infection area (Ar) of the fixed region (L centimeter away from the initial phage inoculation position) was calculated, and the angle of phage spread *θ* was 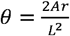. The angle of phage spread θ was still calculated using this formula even when the sides of expansion zones are curved.

### Fluorescence intensity assay

To investigate the location difference of the infected and uninfected cells in the propagating front (Fig. 4E), the plates were scanned by Nikon Ti-E microscopy equipped with a 10× phase contrast objective (Nikon CFI Plan Fluor DL4X F, N.A. 0.13, W.D. 16.4 mm, PhL), the green and red fluorescence was taken with a 2-ms and 500-ms exposure time, respectively. The details were as follows: firstly, as the above-mentioned method, the semi-solid D-RDM medium plate with 0.25% agar and 20 ng/mL aTc was prepared, and the strain ZF1-RFP was cultured; Second, the strain ZF1-RFP was inoculated at the line 2 cm away from the center of the plate and 2 μL 10^9^ pfu/mL M13-GFP phage was inoculated 1 cm away from the bacteria position, and the plate was incubated at 37°C for 10 hours; Third, the Fluorescence intensity of the whole propagating front in the plate was scanned by the Nikon Ti-E microscopy and the data was exported by the NIS-Elements AR software (ver. 4.50.00). The red fluorescence intensity represents the host cell ZF1-RFP including the uninfected and infective cells (infected cells + recovered cells), and the green fluorescence intensity represent the infective cells which express the green fluorescence gene of M13-GFP because of the phage infection. In the back of the infected zone, both fluorescent signals could be normalized by assuming the majority of the cells had been infected (38). Hence, the uninfected cell can be represented by the red fluorescence intensity minus the green fluorescence intensity. This was processed using a custom-written MATLAB code.

## Acknowledgments

The authors thank C. Liu for sharing *E. coli* strain, Y. Bai, S. Huang for discussions, comments, and technical support. This work is partially supported by the National Key Research and Development Program of China (2021YFA0911100), Strategic Priority Research Program of the Chinese Academy of Sciences (XDB0480000), the National Natural Science Foundation of China (32301223,32071417, 32025022).

## Data availability

Major experimental data supporting the findings of this study are available within the main text and Supplementary Information. Modeling and analysis code have been submitted to GitHub with a link https://github.com/YiZhangsiat/Host-Phage-Expansion.

