## Supplementary material for "Navigated range expansion promotes migratory culling": Fig. S,

Xiongfei Fu

**This PDF file includes:**

Supporting text

Figures S1 to S21

Tables S1 to S4

Legends for Movies S1 to S2

SI References

**Other supporting materials for this manuscript include the following:**

Movies S1 to S2

Supporting Information Text

To understand the dynamics of bacteria-phage co-propagation, we extended a growth-expansion model that incorporates bacterial chemotaxis and M13 phage infection kinetics. The model provided a comprehensive picture of the formation dynamics of the fan-shaped infected zone, recaptured the experimental results, and enabled us to examine the migratory culling mechanism resulting from the spatial sorting structure between uninfected and infected cells in the co-propagating front. In this section, we describe in detail the model derivation and its dependence on the functional form and parameter values.

In Section 1, we modified the Susceptible Infected Recovered kinetics (SIR) model to describe the dynamics of bacteria and bacteriophage by incorporating external nutritional restrictions and the influence of viral amplification. We then examined the effects of various mathematical formulations of phage adsorption, bacterial growth rates, and phage infection on modeling bacteria-phage interactions.

In Section 2, we integrated bacterial navigated range expansion with this modified SIR model to develop a Range Expansion with Susceptible Infected Recovered kinetics (RESIR) model. In this model, bacterial motility and chemotaxis are represented by the Keller-Segel model. Additionally, we investigated the effects of phage diffusion, phage infection modes, phage adsorption, and bacterial growth burden from phage infection.

1. **Susceptible Infected Recovered kinetics (SIR) model in liquid culture**
   1.1 **Derivation of the Modified SIR model**

One of the classical models for modeling the spread of a viral infection is the SIR model (1–4), which tracks the fraction of a population in three groups: susceptible individuals *S*, infected individuals *I*, and recovered individuals *R*. Susceptible individuals can become infected after contacting the virus, and infected individuals can recover after developing immunity. This model has been used to gain important insights into the progression of new epidemics in an idealized susceptible population with random mixing.

In contrast to many other phages, M13 phage infection is characterized as chronic. Specifically, infected host cells are not killed; instead, progeny phages are continuously produced and extruded through the cell membrane as the infected cells continue to grow at a slower rate (5–8). The production of phages by infected bacterial cells continues until the cells recover from the phage infection (Fig. S1). Once recovered, the cells grow as fast as susceptible ones (Fig. S1C) and produce progeny phages at a much lower level compared to freshly infected cells (7–9). Additionally, recovered host cells cannot be re-infected by free phage particles (9). These unique features make it suitable to describe the M13 phage infection process with a tailored SIR model.

To gain quantitative insights into bacteria-phage interactions, we modified the SIR model to consider the dynamics of bacterial cell density (susceptible bacteria *S*, infected bacteria *I*, and recovered bacteria *R*), the concentration of the main nutrient (*n*), and the number of M13 phage particles (*p*). The corresponding equations are described as follows:

$\frac{\partial S}{\partial t}=\lambda\left( n \right)S-\kappa_{0}\lambda\left( n \right)pS$	(1)

$\frac{\partial I}{\partial t}=\eta\lambda\left( n \right)I+\kappa_{0}\lambda\left( n \right)pS-\psi I$	(2)

$\frac{\partial R}{\partial t}=\beta\lambda\left( n \right)R+\psi I$	(3)

$\frac{\partial n}{\partial t}=-{\lambda\left( n \right)\left( S+I+R \right)}/{Y_{n}}$	(4)

$\frac{\partial p}{\partial t}=\zeta\left[ \left( 1-\eta\right)\lambda\left( n \right)I+\left( 1-\beta\right)\lambda\left( n \right)R \right]-\varphi S$	(5)

Herein, $\eta$ and $\beta$ represent the growth reduction ratios of infected and recovered bacteria, respectively. $\kappa_{0}$ denotes the phage infection efficiency, $\psi$ is the recovery rate of infected bacteria, $\zeta$ is the phage production rate, $\varphi$ is the phage adsorption rate, $Y_{n}$ is the yield of nutrient consumption.

Bacterial cell growth is described considering the growth limitations caused by nutrient availability and follows a Monod relation (10):

$\lambda\left( n \right)=\frac{\lambda_{0}n}{n+n_{k}}$	(6)

where $n_{k}$ is the Monod constant and $\lambda_{0}$ is the maximum growth rate.

The phage infection term applied to susceptible cells is assumed to follow the cell growth rate, $\lambda\left( n \right)$, reflecting the chronic nature of M13 phage infection. The study of F-pilus production indicates that the total production of filaments increases rapidly during the log phase of growth and drops off and finally ceases as growth of the culture slows down and the culture enters the stationary phase(11). This indirectly suggests that stationary-phase bacteria are less susceptible to infection. In addition, phage infection and reproduction utilize the host cell’s machinery, including replication, transcription, and translation processes, which require external resources (e.g., nutrients) to function. Unlike lytic phages, M13 phage infection does not lead to cell lysis, preventing resource recycling. Therefore, when nutrients are depleted (e.g. $n\to0$, $\lambda\left( n \right)\to0$), we assume phage infection ceases as cell growth halts.

Experimental evidence supports this assumption: we observed that the fan-shaped infected zone remained stable for days until the agar plate dried up (Fig. S2). The boundary between infected and uninfected zones did not expand once formed, as no phage particles or infected cells were detected outside the fan-shaped zone more than 48 hours after its formation. In contrast, using lytic phages (e.g., T7 phage), the infected zones continuously expanded until the agar plate dried up. This phenomenon suggests that the phage cannot infect susceptible cells in the stationary phase.

1.2 **Analysis of the Modified SIR model**

To determine the kinetic parameters of phage infection, we experimentally characterized the infection of M13 phages in a well-mixed batch culture environment (Fig. S1). The growth curves of susceptible bacteria, bacteria in the presence of M13 phages, and recovered bacteria (prepared by culturing FM15 cells overnight with M13 phages) show that M13 phage infection results in temporary growth suppression immediately after infection. This is supported by comparing the growth rates of susceptible and recovered bacteria, with the growth rate of recovered bacteria being almost identical to that of susceptible bacteria (Fig. S1C).

The modified SIR model recaptures the experimental infection kinetics of total bacterial growth and phage titers for various initial phage conditions (Fig. S1). The simulation results demonstrated that the growth of log-phase *E. coli* cells stagnated after phage infection and then recovered to normal after a few minutes. The stagnation of cell growth occurred earlier when the initial phage titer was high (Fig. S1A).

We then explored the effect of phage adsorption in the SIR model. M13 phage is an F-specific phage, and its reproductive life cycle begins with the attachment of the virus to the tip of the F-pili of *E. coli* cells. The F-pilus retracts after virus attachment, pulling the phage closer to the bacterial cell and binding to a coreceptor, after which the phage injects its single-stranded DNA into the bacterial cell. This phage adsorption process was incorporated into the present SIR model.

In the following, we briefly explored the effects of omitting phage adsorption in the SIR model. The equation (5) can then be rewritten as:

$\frac{\partial p}{\partial t}=\zeta\left[ \left( 1-\eta\right)\lambda\left( n \right)I+\left( 1-\beta\right)\lambda\left( n \right)R \right]$	(7)

Combining equations (1)-(4) and (6)-(7), the simplified SIR model can still capture the experimental infection kinetics of total bacterial growth and phage titers for various initial phage conditions (Fig. S16A). Our interpretation of these results is that, although considering the phage adsorption term can result in a lower phage concentration in the environment, it does not fundamentally affect the bacteria-phage interaction in the model. The absorption effect only plays a role when phage concentration is low.

In the above, we assumed bacterial growth followed a Monod relation, which describes the growth rate of bacteria as a function of substrate (nutrient) concentration, capturing the idea that growth increases with substrate availability but eventually plateaus at high concentrations. However, the relationship between growth rate and nutrients is more complex, and the Hill function is also used in biology to model bacterial population growth (12, 13), which can describe cooperative effects with a steep response (controlled by the Hill coefficient). Therefore, we also explored the effects of the bacterial growth function on the behavior of the SIR model, with a Hill coefficient of 2 and as a function of nutrient concentration. Specifically, the equation (6) was written as:

$\lambda(n)=\frac{\lambda_{0}n^{2}}{n^{2}+{n_{k1}}^{2}}$	(8)

Combining the equations (1)-(5) and (8), this model can also capture the experimental infection kinetics of total bacterial growth and phage titers for various initial phage conditions (Fig. S16B).

Furthermore, according to the literature, the phage infection rate can decrease at higher bacterial densities (5). Therefore, a nonlinear form of infection rate can be introduced when constructing a mathematical model of bacteria-phage interaction(14). Similarly, we assumed the phage infection efficiency follows a Hill function:

$\kappa(p)=\frac{\kappa_{0}p^{2}}{p^{2}+{p_{k}}^{2}}$	(9)

Hence, the phage infection efficiency $\kappa_{0}$ in the equations (1)&(2) were written as the form of equation (9). Combining the equations (1)-(5) and (8)-(9), the model can also capture the experimental infection kinetics of total bacterial growth as well as the phage titers for various initial phage conditions (Fig. S16C).

Some experimental studies have observed that an infected cell loses M13 gene at a small rate(15) (the phage was lost about once in six bacterial divisions (16)). We then discussed the effect of phage loss on the interaction of bacteria and phage. The equations (1)&(3) were written as:

$\frac{\partial S}{\partial t}=\lambda\left( n \right)S-\kappa_{0}\lambda\left( n \right)pS+\xi R$	(10)

$\frac{\partial R}{\partial t}=\beta\lambda\left( n \right)R+\psi I-\xi R$	(11)

Herein, $\xi$ is the rate of phage loss. Combining the equations (2), (4)-(6) and (10)-(11), the model can also capture the experimental infection kinetics of total bacterial growth as well as the phage titers for various initial phage conditions (Fig. S16D).

In the above simulations, it is noted that the model parameters $n_{k1}$=5mM, $p_{k}$=0.05 a.u. and $\xi$=7×10^-5^ s^-1^.

In summary, we created a modified SIR model to study bacteria and bacteriophage interactions, including factors like nutrient availability and viral amplification. This model helped us understand how bacteria and phages spread and interact. Our simulations matched experimental data on bacterial growth and phage levels. We found that even without considering phage adsorption, the model still worked well. We also tested different ways to describe bacterial growth, phage infection, and phage loss, showing the model’s flexibility. Overall, our modified SIR model is a strong tool for studying bacteria-phage dynamics, offering flexibility in how we describe these interactions.

2. **Range Expansion with Susceptible Infected Recovered kinetics (RESIR) model**
   2.1 **Derivation of the RESIR model**

Agar, a seaweed extract, is a solidifying agent used in preparing microbiological culture media and has been a standard support matrix for microbial experiments for over a century(17). The effective pore size can be affected by agar gel concentration, as estimated from Ferguson plots(18, 19). In this study, the agar concentration ranged from 0.2% to 0.4%, with an estimated effective pore size of 200 nm to 400 nm.(20). For *E. coli* grown on a semi-solid agar plate, which allows the bacterial cells to swim, the spatiotemporal dynamics are governed by two elements: cell motility (diffusion and chemotaxis) and cell growth. However, the M13 wildtype phage is a flexible thread about 900 nm in length and 7 nm in diameter, meaning that once a progeny phage is produced in a semi-solid agar plate, it is likely to be trapped in its birthplace and cannot move. Additionally, the diffusion coefficient of a similar filamentous bacteriophage in a liquid medium is only 2 μm²/s (21), which is two orders of magnitude smaller than the diffusion coefficient of the bacterial host(22, 23). Therefore, in this study, we ignored phage diffusion when modeling the spatial bacteria-phage interaction.

Based on these assumptions, we developed a RESIR model to gain quantitative insights into the spatial bacteria-phage co-propagation process. This model is based on a navigated range expansion model of bacterial populations(10) and the aforementioned modified SIR model of phage infection. The motility and chemotaxis of bacteria are represented by Keller-Segel-type diffusion and advective terms widely used in the literature(10, 22, 24). The spatiotemporal dynamics of bacterial cell density (susceptible bacteria *S*, infected bacteria *I*, recovered bacteria *R*), concentrations of the main nutrient (*n*) and chemoattractant (*a*), and the number of M13 phage particles (*p*) are described as follows:

$\frac{\partial S}{\partial t}=\mu\nabla^{2}S-\chi\nabla\cdot\left( S\nabla f \right)+\lambda\left( n \right)S-\kappa_{0}\lambda\left( n \right)pS$	(12)

$\frac{\partial I}{\partial t}=\mu\nabla^{2}I-\chi\nabla\cdot\left( I\nabla f \right)+\eta_{1}\lambda\left( n \right)I+\kappa_{0}\lambda\left( n \right)pS-\psi I$	(13)

$\frac{\partial R}{\partial t}=\mu\nabla^{2}R-\chi\nabla\cdot\left( R\nabla f \right)+\beta_{1}\lambda\left( n \right)R+\psi I$	(14)

$\frac{\partial n}{\partial t}=D_{n}\nabla^{2}n-{\lambda\left( n \right)\left( S+I+R \right)}/{Y_{n}}$	(15)

$\frac{\partial a}{\partial t}=D_{a}\nabla^{2}a-\gamma\left( a \right)\left( S+I+R \right)$	(16)

$\frac{\partial p}{\partial t}=\zeta\left[ \left( 1-\eta\right)\lambda\left( n \right)I+\left( 1-\beta\right)\lambda\left( n \right)R \right]-\varphi S$	(17)

Here, $\mu$ and $\chi$ are the effective diffusion and chemotactic coefficients of bacteria, respectively; $D_{n}$ and $D_{a}$ are the diffusion coefficients of the nutrient and chemoattractant, respectively.

The chemotactic movement of bacteria depends on the concentration of the local attractant as follows(25):

$f=log\frac{1+\frac{a}{K_{1}}}{1+\frac{a}{K_{2}}}$	(18)

where $K_{1}$ and $K_{2}$ are the lower and upper Weber offset of attractant sensing.

Bacterial cell growth is described as equation (6) in the above model. Similar as nutrient consumption, chemoattractant uptake is also described by a Monod relation (9, 10):

$\gamma\left( s \right)=\frac{g_{0}a}{a+a_{k}}$	(19)

with the Monod constant $a_{k}$ and the uptake rate of chemoattractant $g_{0}$.

It must be noted that when the chemotactic coefficient $\chi=0$ μm^2^ s^-1^, the above equations (12)-(17) describe the interaction of bacteria and phage under the unguided range expansion. The results shown in Fig. S5 and Fig. S6 were calculated using this model, with the parameters given in Table S4.

In the following, we define boundary and initial conditions and provide the parameters used in this study. All model parameters are summarized in supplementary Table S4.

2.2 **Simulation of the RESIR model**

The equations were integrated in Matlab using second-order centered differences for the spatial derivatives (mesh size 250 µm) and an explicit fourth-order Runge-Kutta routine for temporal integration (time step 10 s) in the Cartesian coordinate system.

**Boundary conditions:**

For cells, nutrients, chemoattractant, and phage, boundary conditions are set to obey zero diffusive flux at the edges of the simulated area:

$\partial_{x}\Phi=0,\text{ }\partial_{y}\Phi=0,\text{ }(\Phi=S,I,R,n,a,p)$.

By default, the habitat size is 20×20 cm^2^.

**Initial conditions:**

**Normal Range Expansion:** The initial susceptible cell density at the center of the simulation area is given by:

$b_{init}=S_{0}e^{-\frac{r^{2}}{r_{0}^{2}}}$ (S0=0.2 OD_600_, and r_0_=1mm),

meaning the simulation starts with a locally restricted susceptible cell density within a radial distance (r < 2 mm) from the center of the simulation region.

**Plane Wave Range Expansion:** The initial susceptible cell density is set at a line 2 cm away from the boundary:

$b_{init}=S_{0}$ (S_0_=0.2 OD_600_).

**Initial Phage Condition:** Located 1 cm away from the initial susceptible cell position:

$p_{init}=P_{0}e^{-\frac{r^{2}}{r_{0}^{2}}}$ (P0=0.1 a.u., and r_0_=1 mm).

**Nutrient and Attractant Concentrations:** Initially homogeneous, given by

$n_{0}=30 mM$ and $a_{0}=100 \mu M$.

2.3 **Analysis of the RESIR model**

The governing equations (12)-(17) were used to describe the spatiotemporal dynamics of bacteria. When *E. coli* grew on a semi-solid agar plate, M13 phage was inoculated in front of it, and a substantial fraction of the expanding cells in the front encountered phages and got infected, resulting in a slowdown of their subsequent growth. Progeny phages were then produced and carried forward along the expansion route by infecting neighboring fresh cells. This combination of cell migration and repeated phage infection was expected to result in the formation of a visible fan-shaped infection zone with lower cell density than the uninfected region. The saturated cell density in the fan-shaped region was lower than that in the uninfected region due to nutrient consumption by phage production, resulting in limited availability for supporting bacterial growth and creating a visible low-density region. The development of the fan-shaped pattern was mainly driven by expansion in the radial direction, supplemented by extension in the lateral direction. The co-propagation migration speed and the lateral expansion speed were both positively dependent on host motility, while the ratio of the two speeds determined the angle of phage spread ($\theta$). Our numerical simulations using reasonable parameter values (Table S4) successfully reproduced the observed fan-shaped pattern in the experiments (Fig. 2).

In developing the spatial interaction model of bacteria and phage mentioned above, we initially did not consider phage diffusion because the length of the phage is considerably larger than the effective pore size of the agar used in this study(5, 20). To explore the effects of potential phage diffusion, we further incorporated a diffusion term into the equation (15):

$\frac{\partial p}{\partial t}=D_{p}\nabla^{2}S+\zeta\left[ \left( 1-\eta\right)\lambda\left( n \right)I+\left( 1-\beta\right)\lambda\left( n \right)R \right]-\varphi S$	(20)

The phage diffusion coefficient used was on the order of the diffusivity of phages in liquids (21). The results demonstrated that introducing phage diffusivity does not qualitatively change the main prediction of the model: a faster host expansion migration hinders virus spread during navigated host range expansion (Fig. S17). Therefore, for simplicity, we chose not to consider phage diffusivity in the RESIR model of this study.

In addition, some experimental studies have observed that an infected cell loses M13 gene at a small rate (15, 16). We then discussed the effect of phage loss on the interaction of bacteria and phage. The equations (12)&(14) can be rewritten as:

$\frac{\partial S}{\partial t}=\mu\nabla^{2}S-\chi\nabla\cdot\left( S\nabla f \right)+\lambda\left( n \right)S-\kappa_{0}\lambda\left( n \right)pS+\xi R$	(21)

$\frac{\partial R}{\partial t}=\mu\nabla^{2}R-\chi\nabla\cdot\left( R\nabla f \right)+\beta_{1}\lambda\left( n \right)R+\psi I-\xi R$	(22)

Combined with the equations (13), (15)-(19) and (21)-(22), we also reproduced the observed fan-shaped pattern seen in the experiments, and the phase transition from co-propagation of bacteria and phage to migratory culling still exist (Fig. S21).

Additionally, according to experimental results reported in the literature and our study, other factors could change the specific mathematical forms in the RESIR model. For example, susceptible cells could be infected by two or more phages (Fig. S18); the phage infection rate can also decrease under higher bacterial density(5), necessitating a nonlinear infection term in the mathematical model of bacteria-phage interaction(14). On the other hand, earlier studies have shown that infected bacteria cannot be re-infected by phages(9). Hence, we constructed another form of the RESIR model to investigate the potential effects of these bacteria-phage interactions:

$\frac{\partial S}{\partial t}=\mu\left( \frac{\partial^{2}S}{\partial x^{2}}+\frac{\partial^{2}S}{\partial y^{2}} \right)-\chi\left( \frac{\partial}{\partial x}\left( S\frac{\partial f}{\partial x} \right)+\frac{\partial}{\partial y}\left( S\frac{\partial f}{\partial y} \right) \right)+\lambda(n)S-\kappa(p )\lambda(n)S$	(23)$\frac{\partial I}{\partial t}=\mu\left( \frac{\partial^{2}I}{\partial x^{2}}+\frac{\partial^{2}I}{\partial y^{2}} \right)-\chi\left( \frac{\partial}{\partial x}\left( I\frac{\partial f}{\partial x} \right)+\frac{\partial}{\partial y}\left( I\frac{\partial f}{\partial y} \right) \right)+\eta\lambda(n)I+\kappa(p )\lambda(n)S-\psi I$	(24)$\frac{\partial R}{\partial t}=\mu\left( \frac{\partial^{2}R}{\partial x^{2}}+\frac{\partial^{2}R}{\partial y^{2}} \right)-\chi\left( \frac{\partial}{\partial x}\left( R\frac{\partial f}{\partial x} \right)+\frac{\partial}{\partial y}\left( R\frac{\partial f}{\partial y} \right) \right)+\beta\lambda(n)R+\psi I$	(25)$\frac{\partial n}{\partial t}=D_{n}\left( \frac{\partial^{2}n}{\partial x^{2}}+\frac{\partial^{2}n}{\partial y^{2}} \right)-\frac{\lambda(n)(S+I+R)}{Y_{n}}$	(26)$\frac{\partial a}{\partial t}=D_{a}\left( \frac{\partial^{2}a}{\partial x^{2}}+\frac{\partial^{2}a}{\partial y^{2}} \right)-\gamma(s)(S+I+R)\text{ }$	(27)$\frac{\partial p}{\partial t}=\zeta((1-\eta)\lambda(n)I+(1-\beta)\lambda(n)R)$	(28)

Here, we considered nutrient complexity by assuming the cell growth following a Hill function (12):

$\lambda(n)=\frac{\lambda_{0}n^{2}}{n^{2}+{n_{k1}}^{2}}$	(29)

We also assumed that the phage infection rate decreases at higher bacterial densities, such that the phage infection efficiency follows a Hill function:

$\kappa(p)=\frac{\kappa_{0}p^{2}}{p^{2}+{p_{k}}^{2}}$	(30)

With the same boundary conditions and initial conditions as the previous model, and the parameters $n_{k1}=5mM$, $\kappa_{0}$=2, $p_{k}=0.05 a.u.$, $\zeta$=5-20 a.u. OD_600_^-1^, $\chi$=0-600 μm^2^ s^-1^, our numerical simulations of this model also reproduced the observed fan-shaped pattern seen in the experiments (Fig. S19), resulting in a better agreement with the experimental data (Fig. S6D, S8B).

In summary, the RESIR model was developed to study the spatial co-propagation of bacteria and bacteriophages. This model integrates bacterial motility and chemotaxis, represented by Keller-Segel type diffusion and advective terms, with a modified SIR model of phage infection. Our simulations demonstrated that the model could successfully reproduce the observed fan-shaped infection patterns seen in experiments. The model’s flexibility allows for the incorporation of various factors, such as nutrient-dependent bacterial growth and phage adsorption processes, making it a robust tool for understanding bacteria-phage interactions. Overall, the RESIR model provides significant insights into the complex dynamics of bacteria-phage interactions in a spatial context, offering a valuable framework for further experimental and theoretical studies.

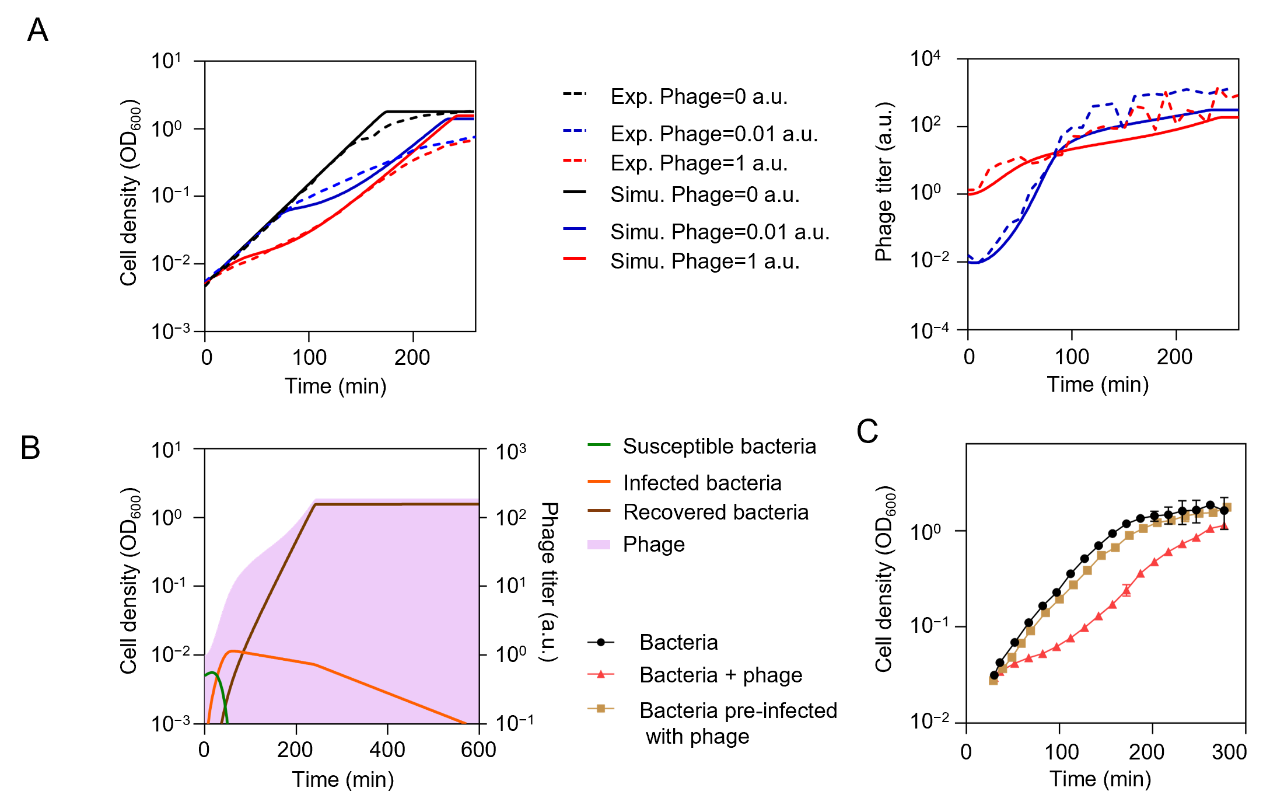

Fig. S1. Effects of phage infection on the growth of E. coli cells in liquid culture. (A), Growth curves of *E. coli* FM15 cells in liquid culture in the presence of phages at different initial titers (left), and growth curves of phages after infection of the host bacteria (right). Bacterial growth clearly slowed down upon phage infection, and gradually recovered to the normal growth rate. Solid and dashed lines represent the simulation and experimental results, respectively. The left y-axis is rescaled to cell density (1 OD_600_=5×10^8^ cells ml^-1^), and the right y-axis to phage titer (1 a.u.=10^9^ pfu ml^-1^). The simulation results are derived from equations (1)-(6) using a phage production rate ζ=700 a.u. OD_600_^-1^, along with the other parameters listed in Table S4. (B), The model simulated the dynamics of bacterial cell density and phage titer with an initial bacterial cell density of 0.005 OD_600_ and a phage titer of 1 a.u. in a well-mixed system. (C), Growth curves of *E. coli* under different conditions of phage infection. *E. coli* FM15 cells were cultured in LB broth. Bacteria are FM15 cells without phage infection; Bacteria + phage are cells with 10^8^ pfu of M13 phages added at the 0-time point. Bacteria pre-infected with phage are cells co-cultured overnight with phages. The pre-infected cells exhibited the same batch growth dynamics as those without phage infection, suggesting that the growth rate of recovered cells resumed.

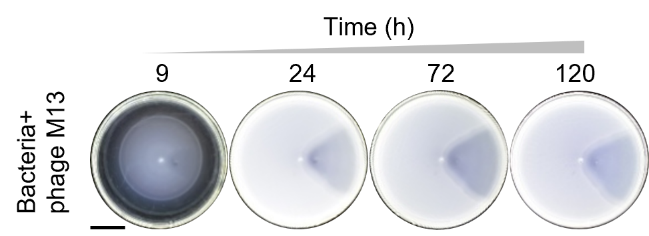

Fig. S2. Time-lapse photographs of typical patterns obtained for bacteria FM15 with phage M13. The scale bar represents 2 cm, and three biological replicates.

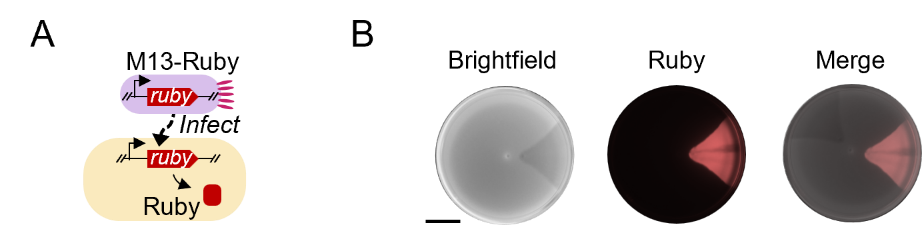

Fig. S3. Formation of the fan-shaped pattern by phage infection during bacterial range expansion. (A), Design of a reporter phage M13-Ruby for fluorescence imaging of phage-infected region. (B), Visualization of infected region. *E. coli* FM15 was inoculated at the center of a semisolid agar plate and the reporter phage M13-Ruby was inoculated 1 cm away from the center. Fluorescence images (Methods) were captured after overnight incubation. The scale bar represents 2 cm. They were repeated independently three times with similar results.

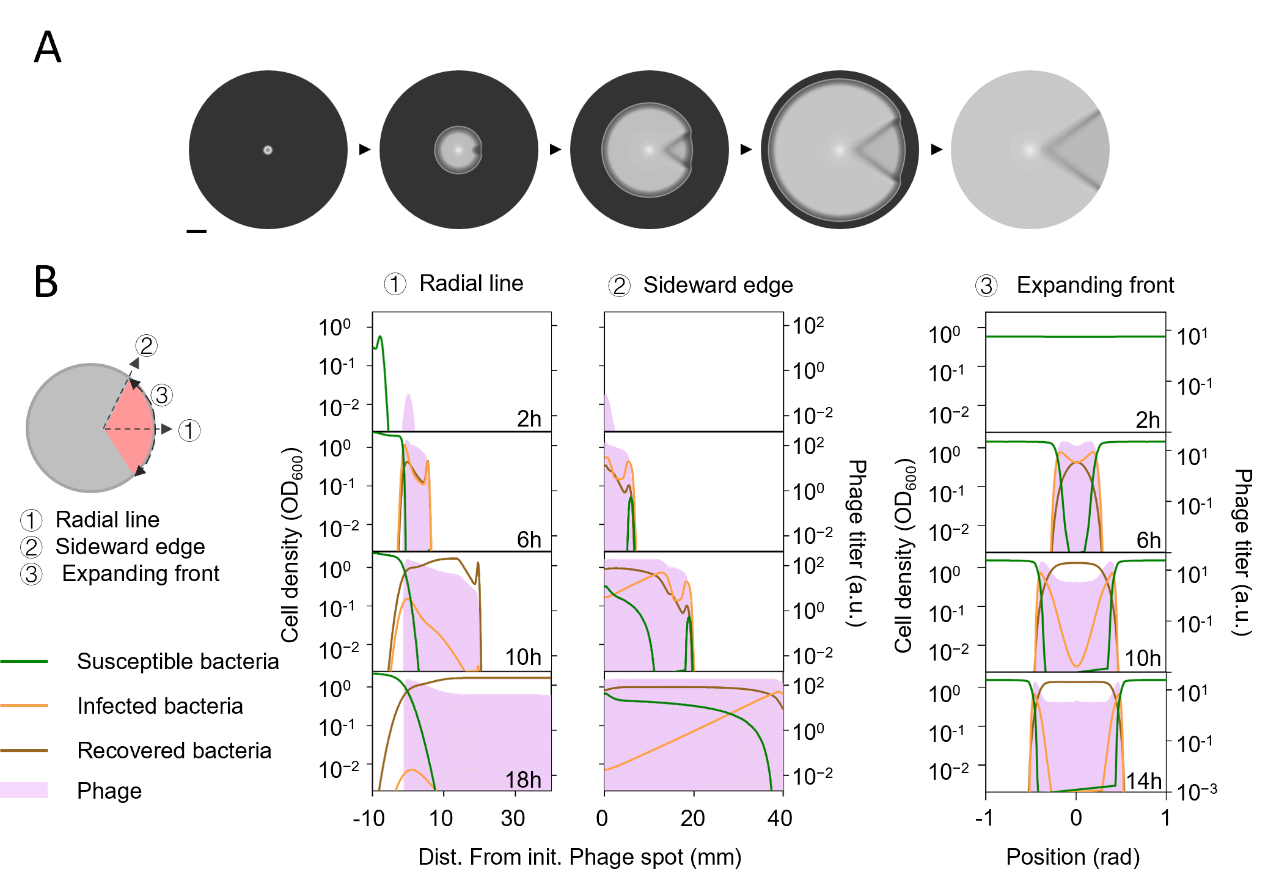

Fig. S4. Kinetic model of the interaction between bacteria and phage. (A) Time-lapsed photographs of typical patterns. scale bar represents 1 cm. (B) Time-lapse plots of the simulated relative bacterial cell density and phage-titer profiles along the radial line ①, the sideward edge ② of the fan-shaped infection region, and the bacteria expanding front ③. The results are obtained from equations (12)-(19) with phage production rate ζ=200 a.u. OD_600_^-1^ and chemotactic coefficient χ=350 μm^2^ s^-1^.

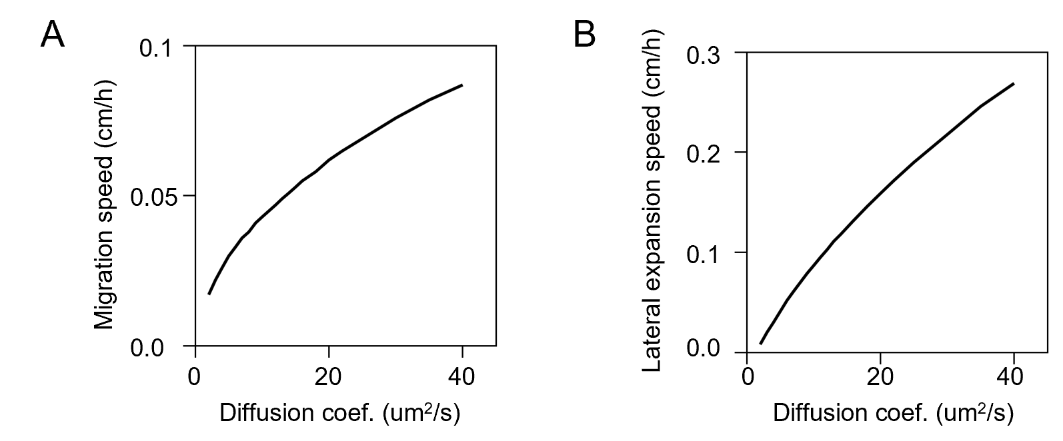

Fig. S5. The simulation of the diffusion coefficient’s effect on expansion speed under the host unguided range expansion (without chemotaxis). (A) The relationship between the host diffusion motility and co-propagation migration speed. (B) The relationship between the host diffusion motility and lateral expansion speed. The results were obtained from equations (23)-(30) with the chemotactic coefficient χ=0 μm^2^ s^-1^ for simulating the host unguided range expansion.

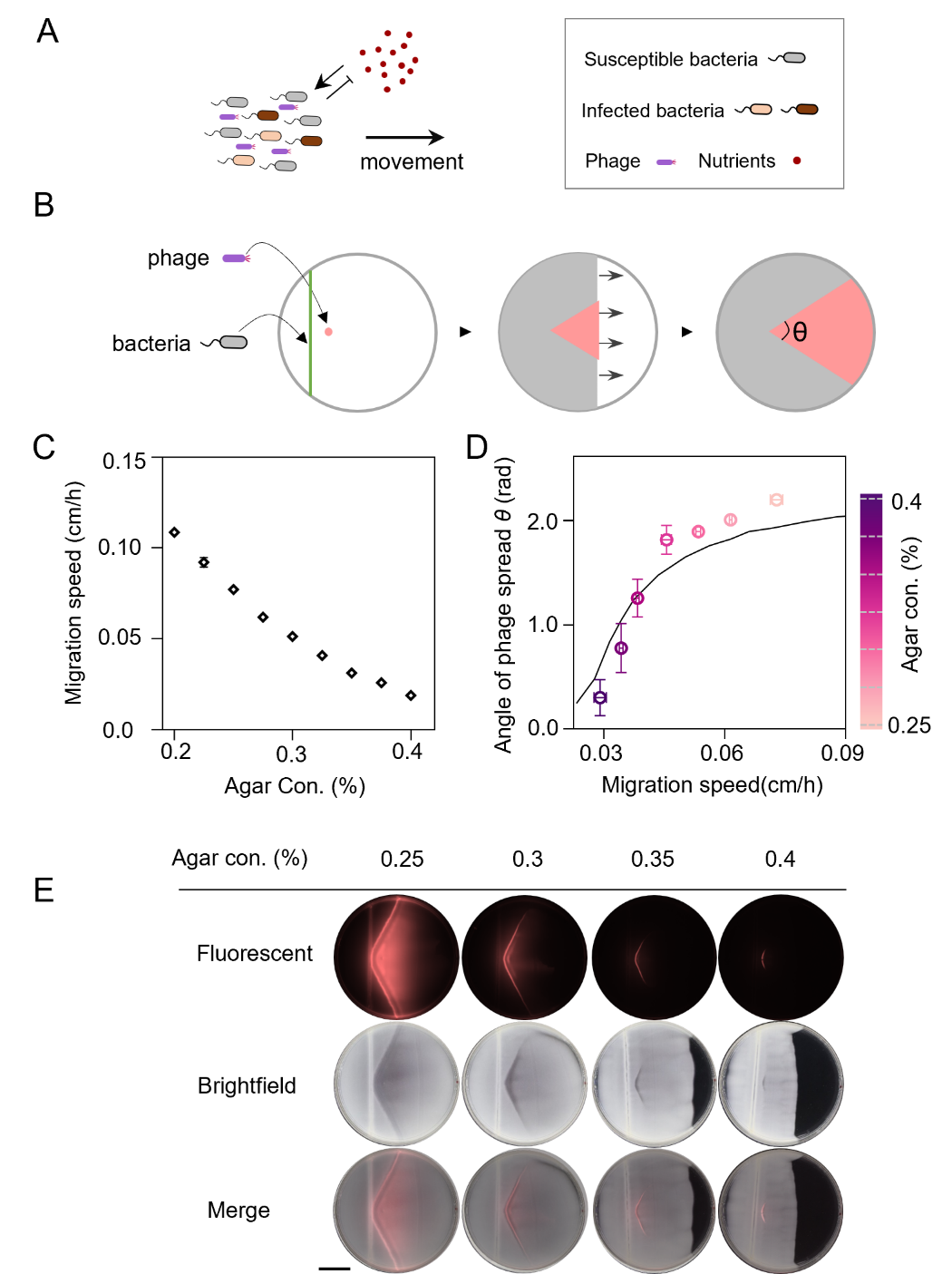

Fig. S6. The phage spread among the bacteria Ft-Δ*cheRcheB* under the plane wave with unguided range expansion. (A) Schematic illustration of the population’s unguided range expansion. (B) Phage propagation during bacterial range expansion under the plane wave leads to the formation of a visible fan-shaped region of lower cell density. (C) The relationship between the agar concentration and the migration speed. Data are mean ± s.d. for n=4 biological replicates. (D) The experiment (circles) and simulation (line) show that the angle of phage spread *θ* is positively correlated with the bacteria migration speed. Data are mean ± s.d. for n=5 biological replicates. The simulation results were obtained from equations (23)-(30) with the chemotactic coefficient χ=0 μm^2^ s^-1^ for simulating the host unguided range expansion. (E) The typical fan-shaped patterns under different agar concentrations. Chemotaxis deficient *E. coli* Ft-Δ*cheRcheB* was inoculated at the line of a semisolid agar plate and the reporter phage M13-Ruby was inoculated 1 cm away from the center. The scale bar represents 2 cm.

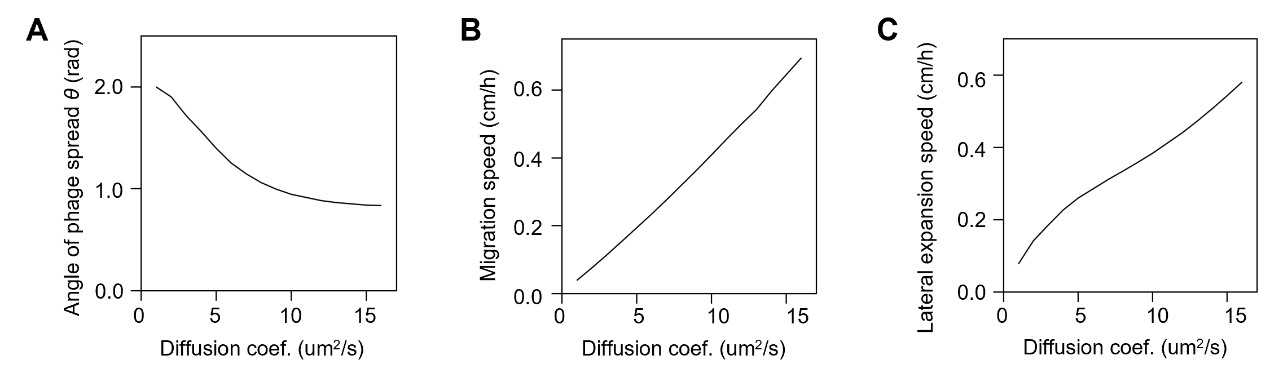

Fig. S7. The simulation of the diffusion coefficient’s effect on expansion speed under the host navigated range expansion. (A) The relationship between the host diffusion motility and the angle of phage spread *θ*. (B) The relationship between the host diffusion motility and co-propagation migration speed. (C) The relationship between the host diffusion motility and lateral expansion speed. The results were obtained from equations (23)-(30) by varying the diffusion coefficients.

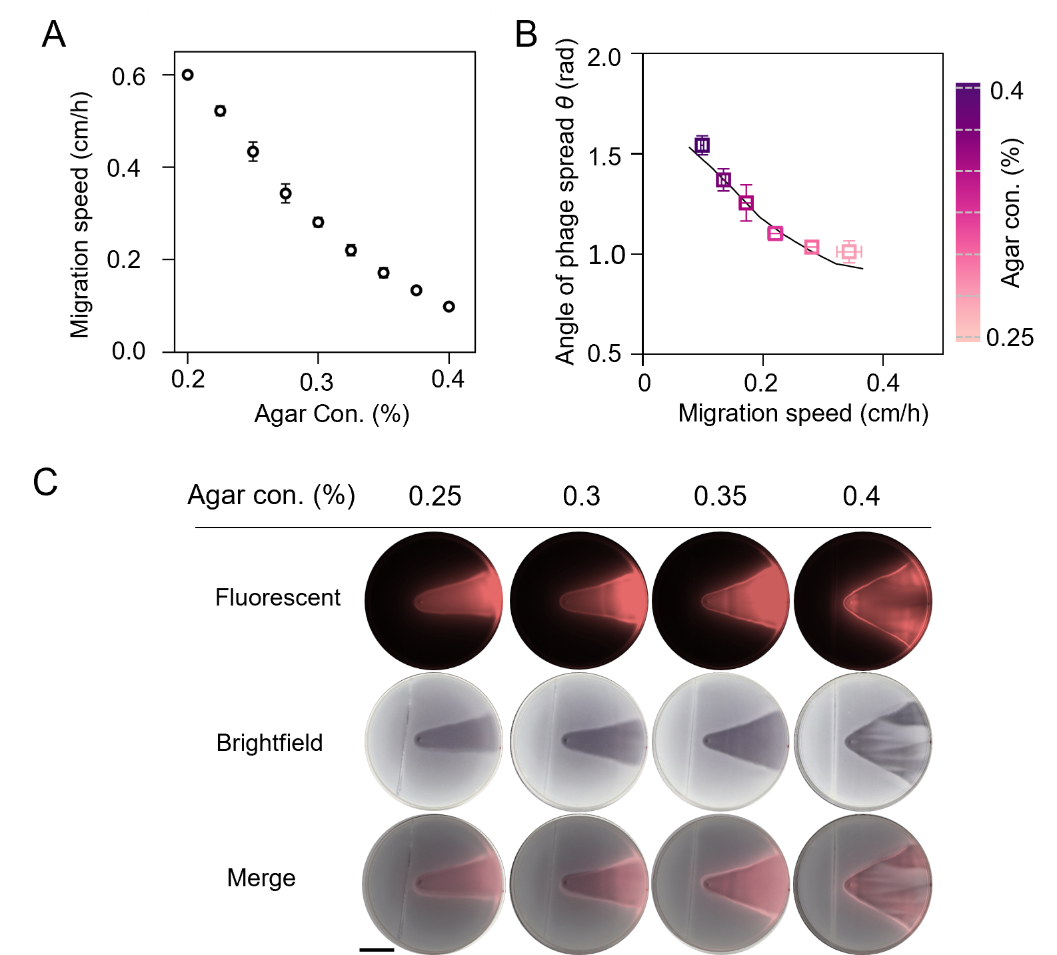

Fig. S8. The phage spread among the bacteria FM15 under the plane wave with navigated range expansion. (A) The relationship between the agar concentration and the migration speed. Data are mean ± s.d. for n=4 biological replicates. (B) The experiment (squares) and simulation (line) show that the angle of phage spread *θ* is negatively correlated with the bacteria migration speed. The simulation results were derived from equations (23)-(30) by varying different diffusion coefficients. Data are mean ± s.d. for n=5 biological replicates. (C) The typical fan-shaped patterns under different agar concentrations. *E. coli* FM15 was inoculated at the line of a semisolid agar plate and the reporter phage M13-Ruby was inoculated 1 cm away from the center. The scale bar represents 2 cm.

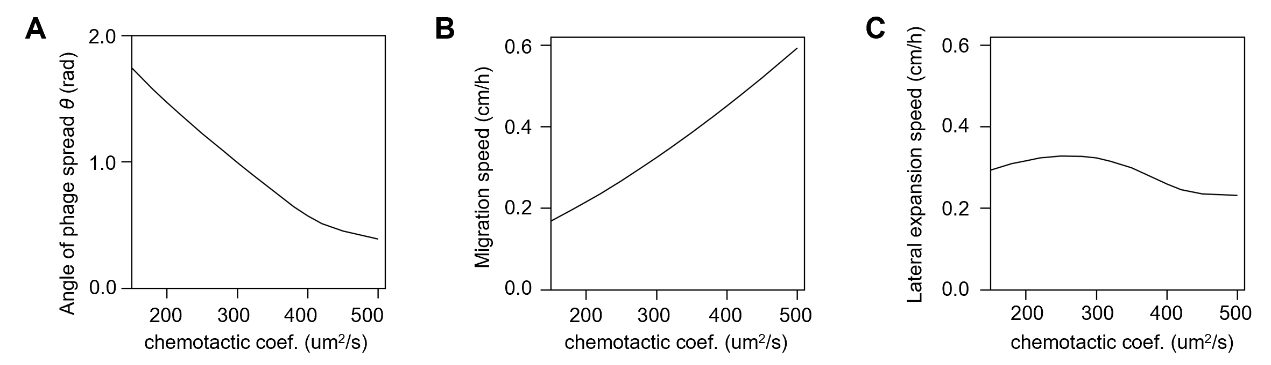

Fig. S9. The simulation of the chemotactic coefficient’s effect on expansion speed. (A) The relationship between the host chemotactic motility and the angle of phage spread *θ*. (B) The relationship between the host chemotactic motility and co-propagation migration speed. (C) The relationship between the host chemotactic motility and lateral expansion speed. The simulation results were obtained from equations (23)-(30) by varying different chemotactic coefficients.

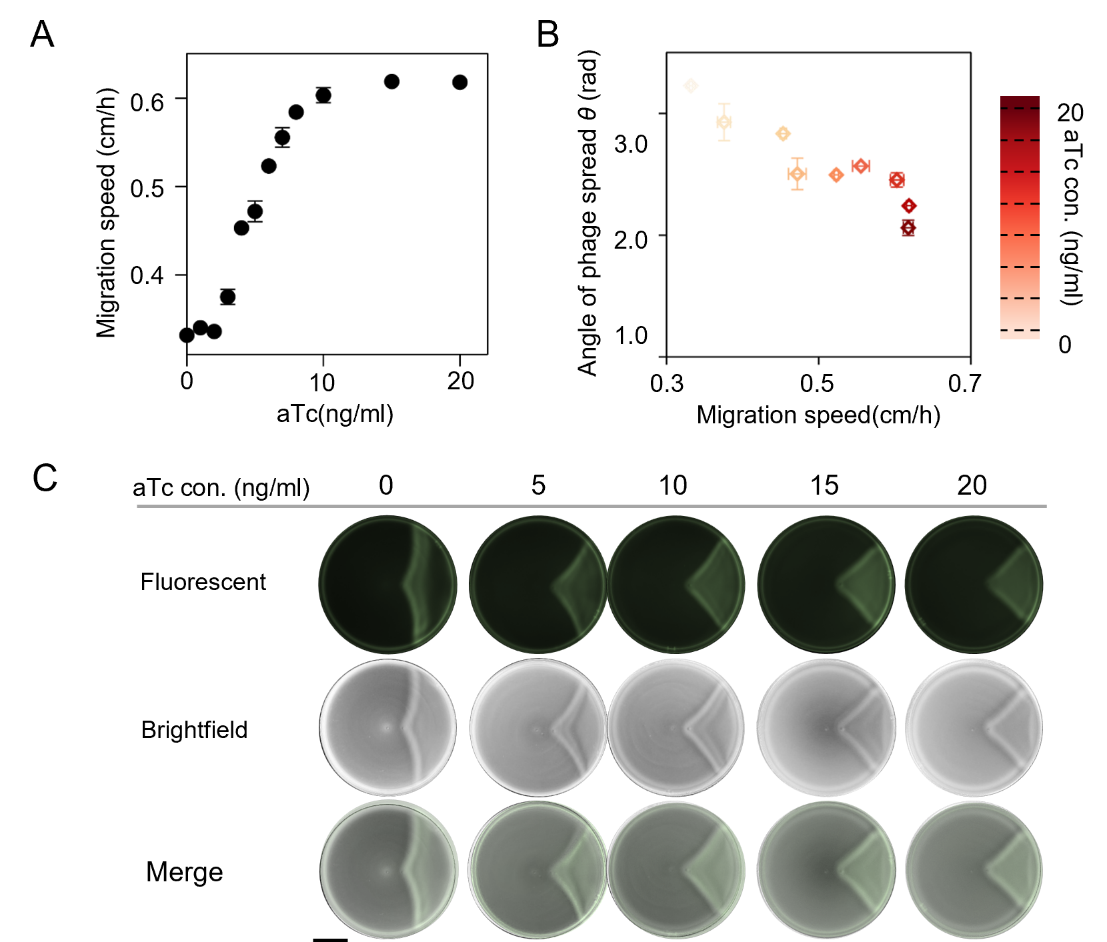

Fig. S10. The phage spread among the bacteria ZF1 regulated by different aTc concentrations. (A) Relationship between the aTc concentration and the migration speed. (B) The experiment shows that the angle of phage spread *θ* is negatively correlated with the bacteria migration speed. (C) The typical fan-shaped patterns under different aTc concentrations. *E. coli* ZF1 was inoculated at the center of a semisolid agar plate and the reporter phage M13-GFP was inoculated 1 cm away from the center. The scale bar represents 2 cm. data are mean ± s.d. for n=5 biological replicates.

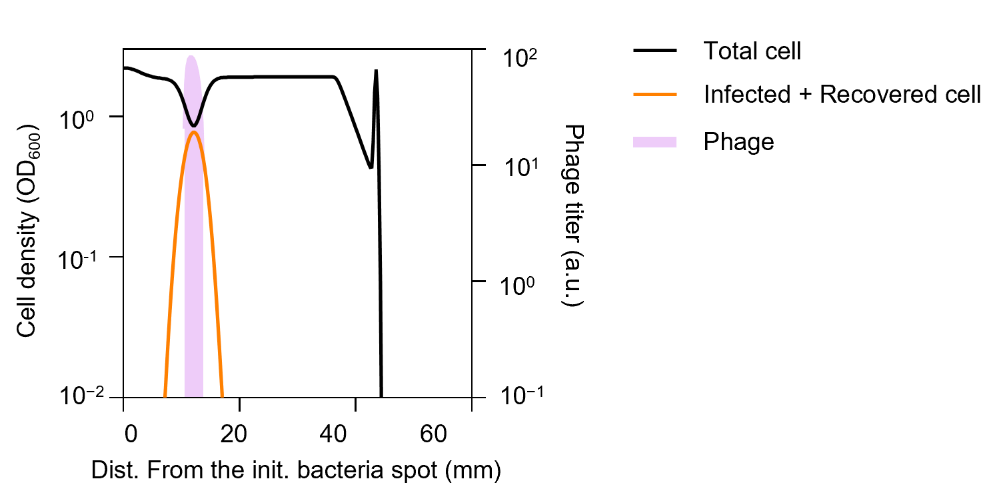

Fig. S11. Profiles of the cell density and phage titer along the central radial line of the fan-shaped infection zone are shown in the simulation results obtained from equations (12)-(19) with a chemotactic coefficient χ=500 μm^2^ s^-1^ and phage production rate ζ=125 a.u. OD_600_^-1^.

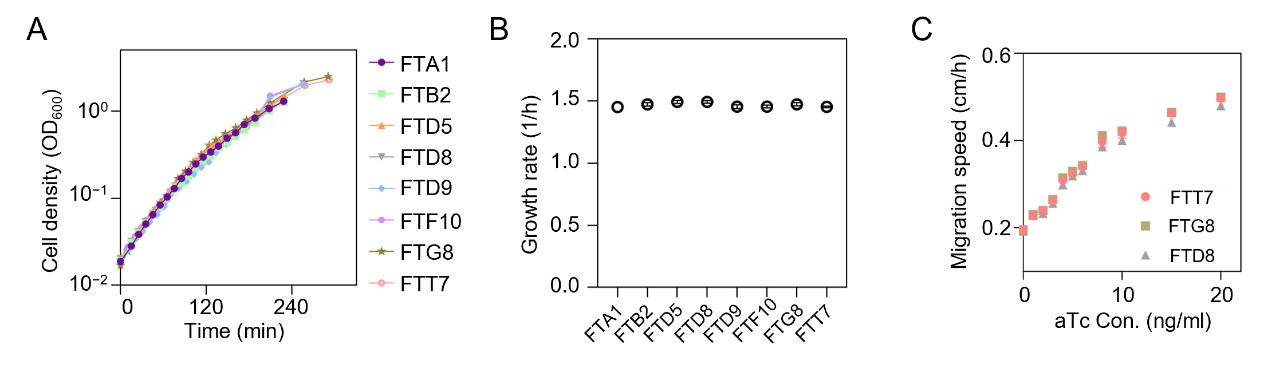

Fig. S12. The phenotypic characteristics of the strains with different T7 RNAP activities. (A) The growth curve for the eight strains with different T7 RNAP activities. (B) The growth rate for the eight strains with different T7 RNAP activities. (C) The relationship between the aTc concentration and the migration speed for the strain FTT7, FTG8 and FTD8. data are mean ± s.d. for n=3 biological replicates.

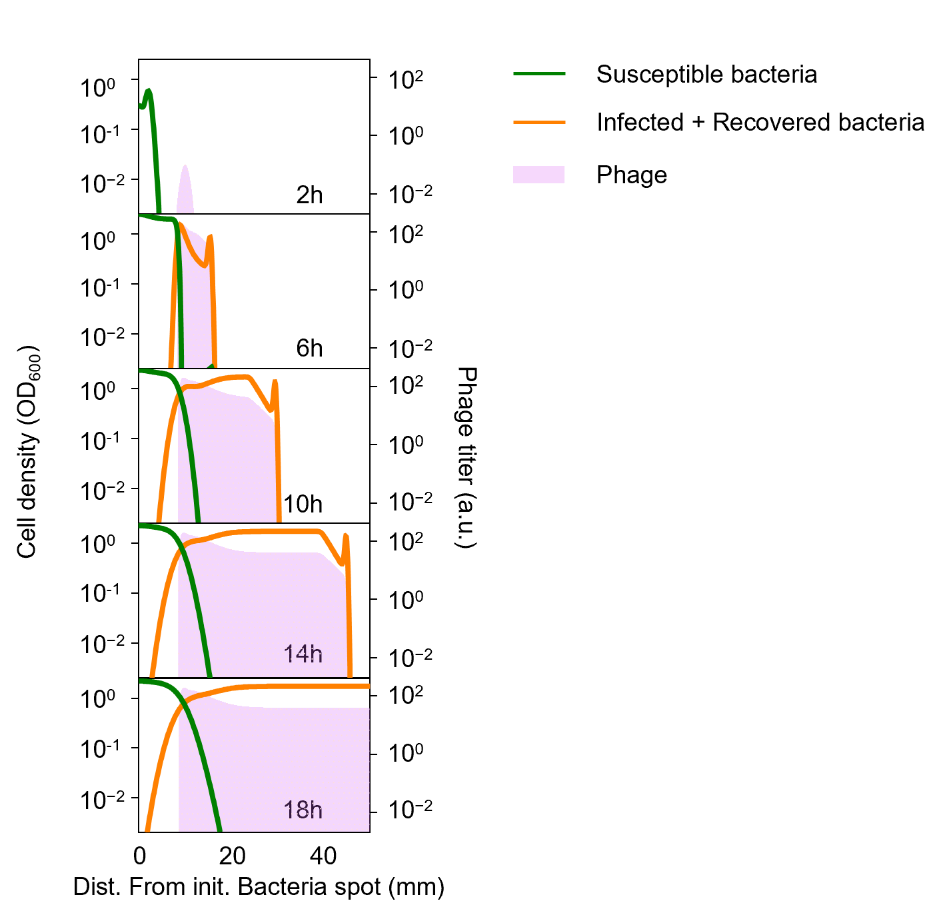

Fig. S13. Time-lapse plots of the simulated relative bacterial cell density and phage profiles under the navigated range expansion along the middle radial axis. The simulation results were obtained from equations (12)-(19) with the parameter chemotactic coefficient χ=350 μm^2^ s^-1^ and phage production rate ζ=200 a.u. OD_600_^-1^.

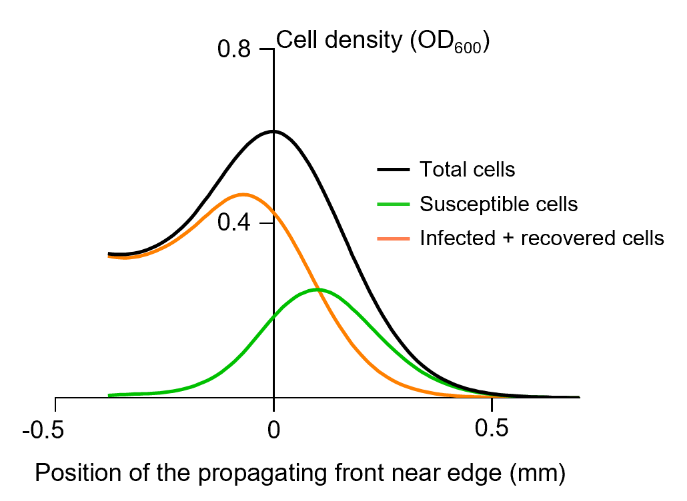

Fig. S14. The simulated density profiles along the leading propagating front near the edge between infected and uninfected zone (Results from equations (12)-(19) with parameter chemotactic coefficient χ=200 μm^2^ s^-1^ and phage production rate ζ=100 a.u. OD_600_^-1^). Black, green, and orange lines represent total, susceptible, infected + recovered bacterial cells, respectively.

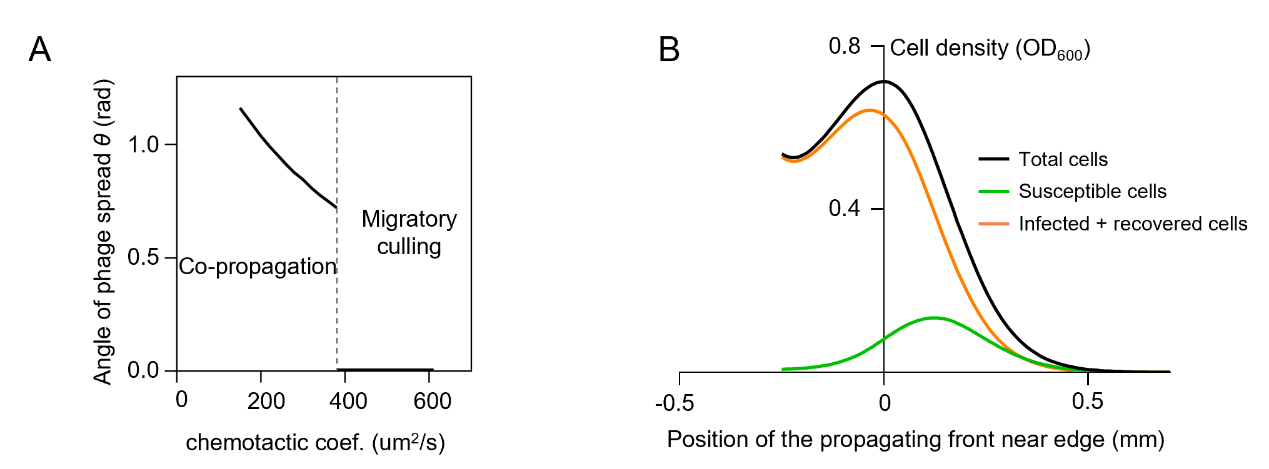

Fig. S15. The simulation results of the RESIR model omitting the bacterial growth burden. (A) The relationship between the host chemotactic motility and the angle of phage spread *θ*. (B) The simulated density profiles along the leading propagating front near the edge between the infected and uninfected zone. The results of (A) and (B) were obtained from equations (12)-(19) with phage production ζ=100 a.u. OD_600_^-1^, chemotactic coefficient χ=200 μm^2^ s^-1^, and the growth reduction ratio of infected and recovered cell η=1, β=1. The results showed that the spatial sorting of uninfected and infected hosts in the navigated propagation front still exists even without the temporary growth reduction of bacteria after phage infection. Black, green, and orange blue lines represent total, susceptible, infected+ recovered bacterial cells, respectively.

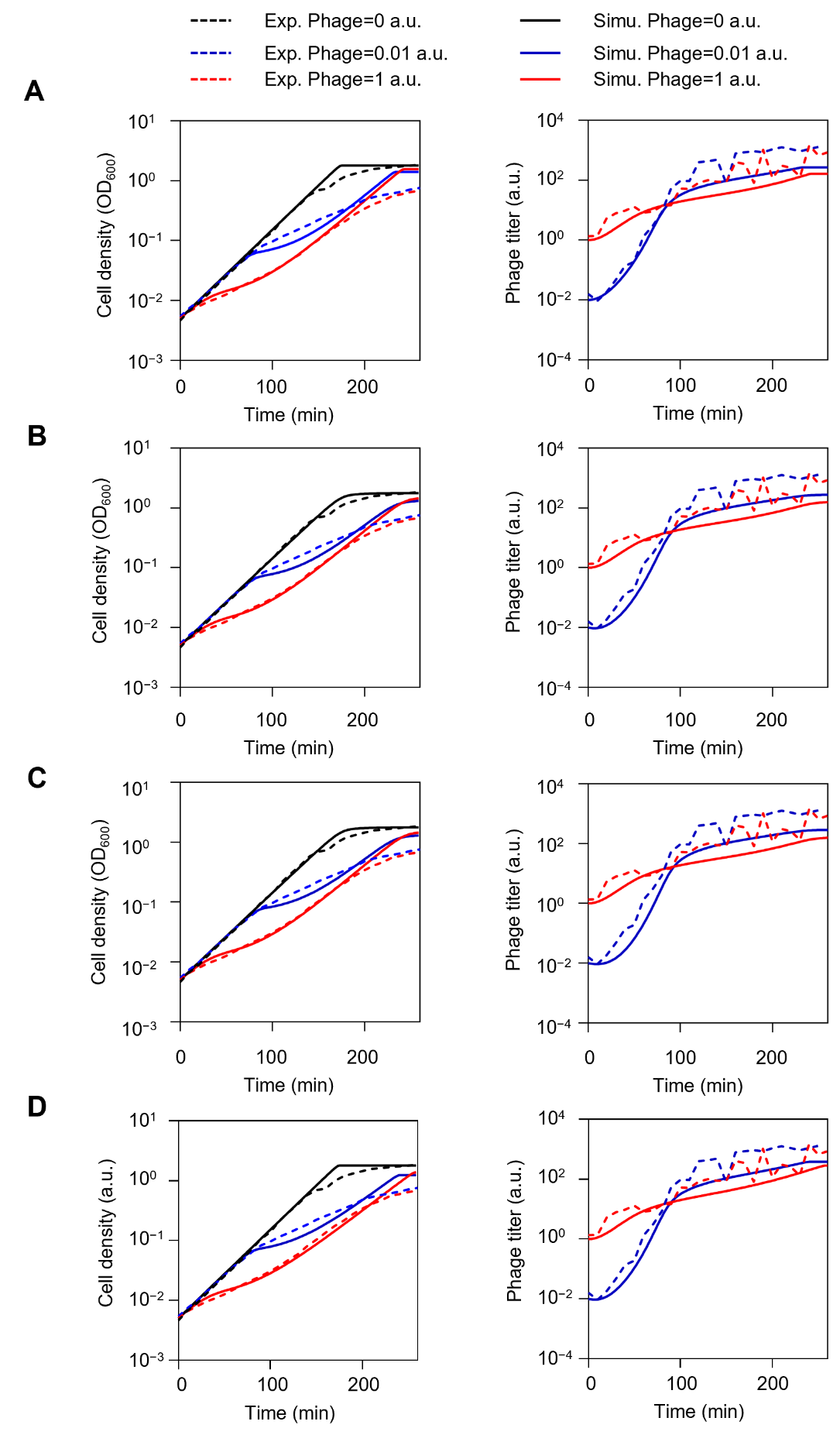

Fig. S16. The simulation of the SIR model under different assumptions. (A) The simulation of SIR model ignoring phage adsorption (obtained from equations (1)-(4)&(6)-(7)). (B) The simulation of SIR model in which the growth rate follows a Hill function (obtained from equations (1)-(5)&(8)). (C) The simulation of SIR model in which the growth rate and phage infection rate both follow a Hill function (obtained from equations (1)-(5)&(8)-(9)). (D) The simulation of SIR model with phage loss (obtained from equations (2), (4)-(6)&(10)-(11)). The solid and dashed lines represent the simulation and experimental results, respectively. These results of different ways to describe bacteria growth, phage infection, and phage loss, show the SIR model’s flexibility.

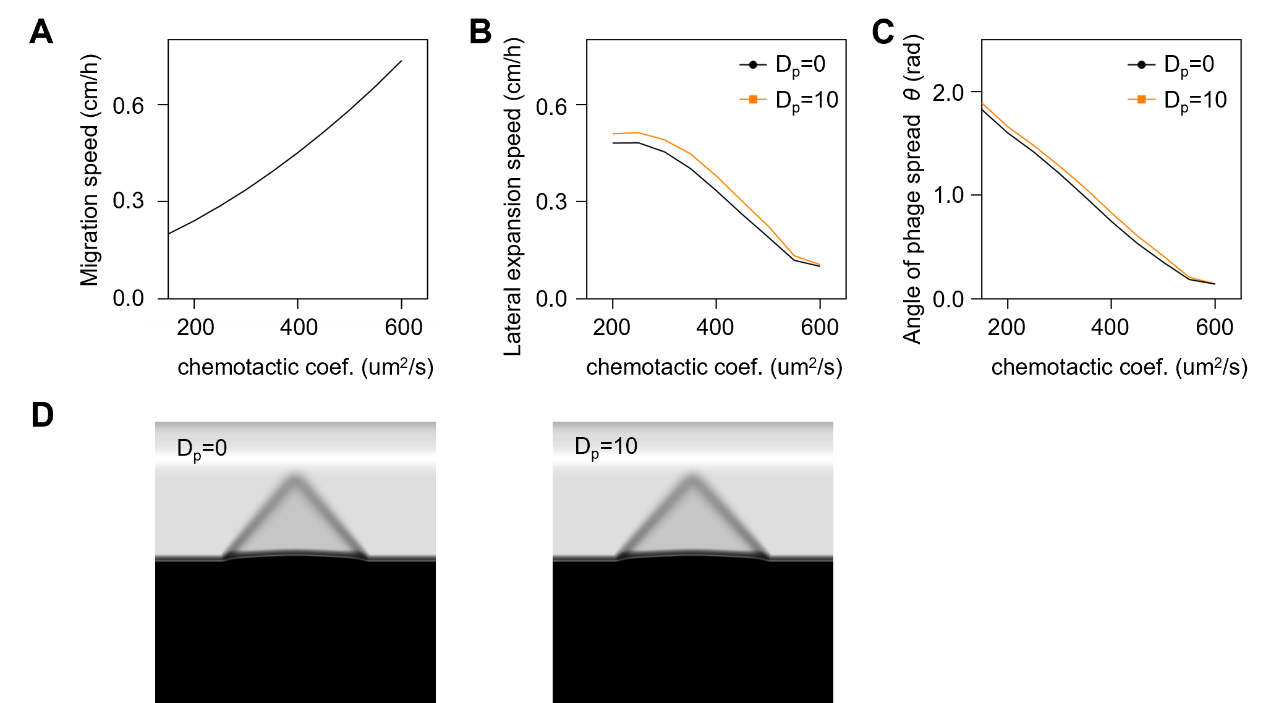

Fig. S17. The simulated effect of the chemotactic coefficient on expansion speed with the RESIR model with/without phage diffusion. (A) The relationship between the host chemotactic motility and co-propagation front speed. (B) The relationship between the host chemotactic motility and lateral expansion speed with varying phage diffusion D_p_. (C) The relationship between the host chemotactic motility and the angle of phage spread *θ* with varying phage diffusion D_p_. (D) The typical fan-shaped patterns under varying agar phage diffusion D_p_. The results were obtained from equations (12)-(16) and (18)-(20) with phage diffusion coefficient D_p_=0 μm^2^ s^-1^ and 10 μm^2^ s^-1^.

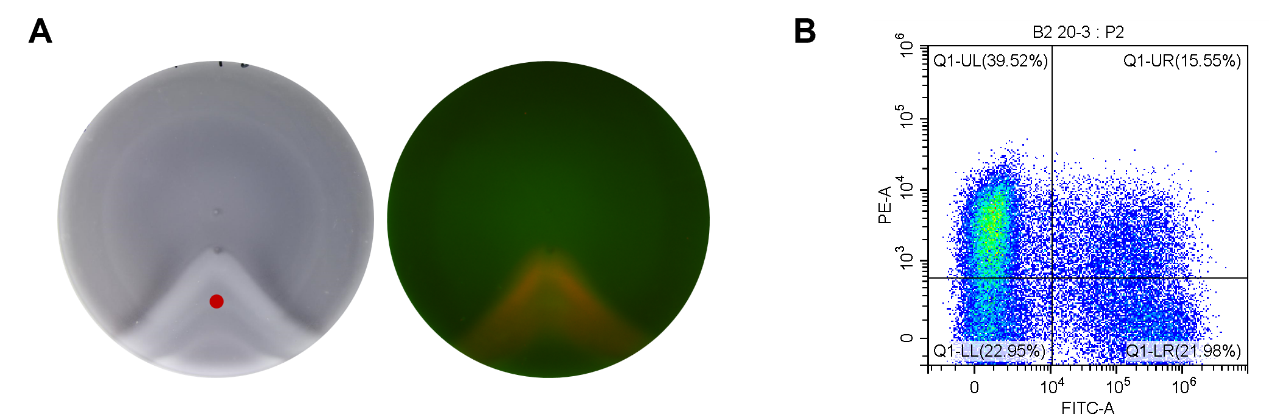

Fig. S18. Measurement of *E.coli* ZF1 infected by two phages simultaneously. (A) Visualization of the infected region. *E. coli* ZF1 was inoculated at the center of a semisolid agar plate and the well-mixed phage M13-GFP:M13-RFP=10^9^:10^9^ pfu/ml was inoculated 1 cm away from the center. Fluorescence images (Methods) were captured after overnight incubation. (B)Four-quadrant diagram obtained for the measurement of phage infection in the red dots as shown in (A) based on flow cytometry.

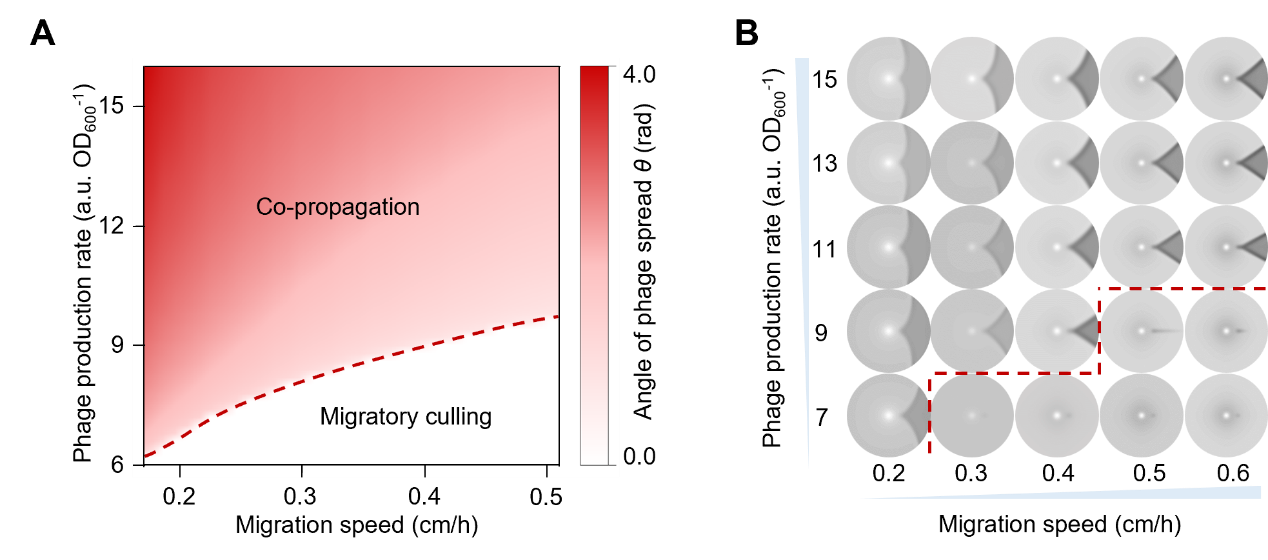

Fig. S19. Prediction of bacterial-viral co-propagation using the RESIR model with equations (23)-(30). (A) the simulated phase transition of bacterial-viral co-propagation. (B) The typical fan-shaped patterns under the different migration speeds and phage production rates.

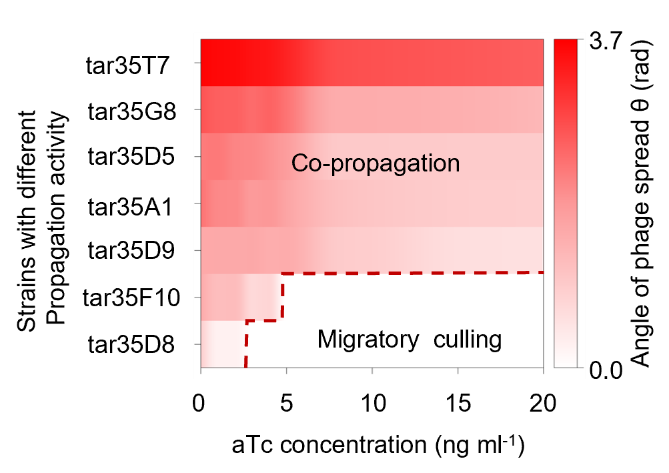

Fig. S20. The repeat experiment of Fig. 4D in main text. The phage spread transition of seven strains with varying phage propagation activity (The numbers on the y axis correspond to the strains with different T7 promoter variants, and the sequences are shown in Supplementary Table S2) under different aTc concentrations. The result verifies that the angle of phage spread *θ* is negatively correlated with the bacteria migration speed and positively correlated with the phage production rate, and the migratory culling exists under the higher migration speed and lower phage production rate.

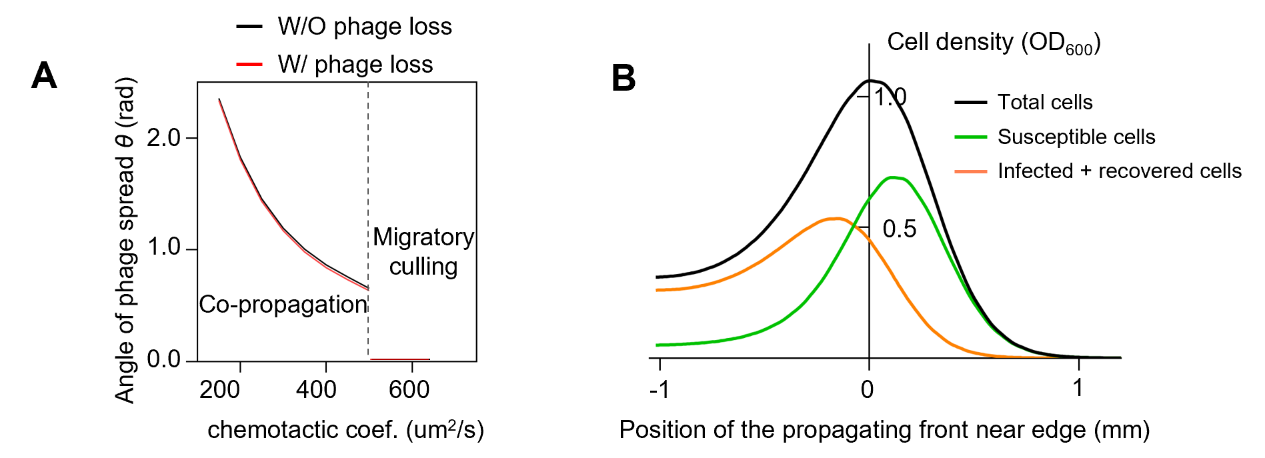

Fig. S21. The simulation results of the RESIR model with phage loss. (A) The relationship between the host chemotactic motility and the angle of phage spread *θ* with and without phage loss. (B) The simulated density profiles along the leading propagating front near the edge between infected and uninfected zone. The results were obtained from equations (13), (15)-(19) and (21)-(22) with phage production ζ=130 a.u. OD_600_^-1^, chemotactic coefficient χ=150-650 μm^2^ s^-1^, Black, green, and orange blue lines represent total, susceptible, infected+ recovered bacterial cells, respectively.

Table S1. Bacteria strains used in this study

| **Strain name** | **Genotype and remarks** | **Reference or source** |
| --- | --- | --- |
| CLM | A motile wild type *E.coli* K12 strain AMB 1655. | Ref.(12) |
| ER2738 | *E.Coli* K12. F plasmid with tetracycline-resistant. | NEB |
| FkP | Δ*tetR*, Δ*tetA* at F plasmid of ER2738,and Kan resistant gene inserted. | This study |
| FM15 | A strain conjugated the *E.coli* CLM and the *E.coli* ER2738 (F plasmid provider) | Ref.(9) |
| Ft-MGΔ*cheRcheB* | Δ*cheR*, Δ*cheB* at FM15 | This study |
| MGT | Δ*tar* at CLM, bla: P_tet_ - *tetR*-*tar* inserted at the attB site of CLM. The expression of *tar* in under the control of a P_tet_ -*tetR* feedback loop. Anhydrotetracycline(aTc) was used as the inducer. | This study |
| ZF1 | Conjugation of the MGT and the FkP (F plasmid with Kan resistant provider) | This study |
| FTT7 | ZF1+ a plasmid T7 with a wild type T7 promoter | This study |
| FTA1/FTD5/FTD8/FTD9/FTF10/FTG8 | ZF1+ a plasmid 4A1/1D5/1D8/4D9/4F10/1G8 with a T7 promotor variant (the promoter sequence is shown in Table S2) | This study |
| ZF1-RFP | ZF1+a plasmid with RFP | This study |

Table S2. Sequences of T7 promoter variants(9)

| **Number** | **Sequence** | **Rel. expression activity of WT T7 RNAP** |
| --- | --- | --- |
| T7 | TAATACGACTCACTATAGGGAGA | 100 |
| 4A1 | TAATACGCCTCACCACTGGGAGA | 0.3379±0.0842 |
| 1D5 | TAGTGCGACTCACAATAGGGAGA | 0.3785±0.0866 |
| 1D8 | TGTATGAACTCACGTTGGGGAGA | 0.0054±0.0048 |
| 4D9 | AAATAAGCCTCACCGTAGGGAGA | 0.1210±0.0496 |
| 4F10 | TAAAGTGGCTCACTATTGGGAGA | 0.025±0.0094 |
| 1G8 | TAATACGCCTCACCACAGGGAGA | 0.7349±0.2127 |

Table S3. Plasmids used in this study

| **Plasmid name** | **Antibiotic resistance** | **Origin of replication** | **Promoter** | **Genes** |
| --- | --- | --- | --- | --- |
| AP-T7 | Spe | puC | P_T7_ | *gIII* |
| AP-A1 | Spe | puC | P_T7_ variant | *gIII* |
| AP-D5 | Spe | puC | P_T7_ variant | *gIII* |
| AP-D8 | Spe | puC | P_T7_ variant | *gIII* |
| AP-D9 | Spe | puC | P_T7_ variant | *gIII* |
| AP-F10 | Spe | puC | P_T7_ variant | *gIII* |
| AP-G8 | Spe | puC | P_T7_ variant | *gIII* |

Table S4. Model parameters

| Parameter | symbol | Value | Reference or parameter variation |
| --- | --- | --- | --- |
| Effective diffusion coefficient of bacteria | μ | 1-60 μm^2^ s^-1^ | Fit to match expansion speed for expansion in reference condition.  See Fig. S6 |
| Chemotactic coefficient of bacteria | χ | 150 - 700 μm^2^ s^-1^ | Fit to match expansion speed for expansion in reference condition.  See Fig. S8 |
| Diffusion coefficient of nutrient | D_n_ | 800 μm^2^ s^-1^ | Ref.(26) |
| Diffusion coefficient of chemoattractant | D_a_ | 800 μm^2^ s^-1^ | Ref.(26) |
| Diffusion of phage | D_p_ | 10 μm^2^ s^-1^ | Ref.(21) |
| Growth reduction ratio of infected cell | η | 0.1 | Fit to match the growth curve of *E. coli* infected by M13 phage. Fig. S1 |
| Growth reduction ratio of recovered cell | β | 0.9 | Fit to match the growth curve of *E. coli* infected by M13 phage. Fig. S1 |
| Recovery rate of infected bacteria | $\psi$ | 10^-4^ s^-1^ | Fit to match the growth curve of *E. coli* infected by M13 phage. Fig. S1 |
| Phage adsorption rate | $\varphi$ | 5×10^-4^ a.u. OD_600_^-1^ s^-1^ | Ref.(5) |
| Phage production rate | $\zeta$ | 70-700 a.u. OD_600_^-1^ | Production rate varied to match the results of fan shape size at different levels of *gIII* expression |
| Growth rate | *λ*_0_ | 1.5 h^-1^ | The average growth rate in D-RDM medium. |
| Monod constant | $n_{k}$ | 0.4 mM | Defined by fitting the growth curve of *E. coli*. Fig. S1 |
| Lower Weber offset attractant sensing | K_1_ | 3.5 μM | Ref.(10, 27) |
| Upper Weber offset attractant sensing | K_2_ | 1000 μM | Ref.(10, 27) |
| Phage infection efficiency | $\kappa_{0}$ | 0.7 a.u.^-1^ | Fit to match the growth curve of *E. coli* infected by M13 bacteriophage. Fig. S1 |
| Uptake rate chemoattractant | g_0_ | 9 uM min^-1^ OD_600_^-1^ | Ref.(22) |
| Monod constant of Attractant | $a_{k}$ | 1 μM | Ref.(28) |
| Yield of nutrient consumption | $Y_{n}$ | 0.064 OD_600_ mM^-1^ | Ref.(10) |

Movie S1 (separate file). A time-lapse movie showing the infection dynamics of the M13-GFP phage. 2 μL of the strain *E. coli* ZF1 (OD_600_=0.2-0.3) was inoculated at the center of a semi-solid agar plate. 2 μL M13-GFP phages with a concentration 10^9^ pfu/ml were inoculated 1 cm away from the bacteria position, and then incubated at 37°C for several hours until the bacteria occupied the whole plate. the plates were scanned by Nikon Ti-E microscopy equipped with a 4× phase contrast objective every 20 min. The movie shows the results after 380 minutes. The culture condition: D-RMD medium + 100 μM aspartate +10 ng mL-1 aTc + 0.25% agar concentration + 10 μg mL-1 kanamycin + 20 μg mL-1 ampicillin.

Movie S1 (separate file). A time-lapse movie showing the infection dynamics of the M13-GFP phage. The repeat experiments of Movie S1

Dataset S1 (separate file). The experimental data used in the figures of this manuscript.

Dataset S2 (separate file). The experimental data used in the figures of SI.

28.	D. Schellenberg, E. Furlongs, Resolution of the Multiplicity of the Glutamate and Aspartate Transport Systems of Escherichia coli.
